# Variational Infinite Heterogeneous Mixture Model for Semi-supervised Clustering of Heart Enhancers

**DOI:** 10.1101/442392

**Authors:** Tahmid F. Mehdi, Gurdeep Singh, Jennifer A. Mitchell, Alan M. Moses

## Abstract

**Motivation:** PMammalian genomes can contain thousands of enhancers but only a subset are actively driving gene expression in a given cellular context. Integrated genomic datasets can be harnessed to predict active enhancers. One challenge in integration of large genomic datasets is the increasing heterogeneity: continuous, binary and discrete features may all be relevant. Coupled with the typically small numbers of training examples, semi-supervised approaches for heterogeneous data are needed; however, current enhancer prediction methods are not designed to handle heterogeneous data in the semi-supervised paradigm.

**Results:** We implemented a Dirichlet Process Heterogeneous Mixture model that infers Gaussian, Bernoulli and Poisson distributions over features. We derived a novel variational inference algorithm to handle semi-supervised learning tasks where certain observations are forced to cluster together. We applied this model to enhancer candidates in mouse heart tissues based on heterogeneous features. We constrained a small number of known active enhancers to appear in the same cluster, and 47 additional regions clustered with them. Many of these are located near heart-specific genes. The model also predicted 1176 active promoters, suggesting that it can discover new enhancers and promoters.

**Availability:** We created the ‘dphmix’ Python package: https://pypi.org/project/dphmix/

## Introduction

Enhancers are cis-regulatory elements in DNA that can influence expression levels of target genes when bound to transcription factors (TFs). They are thought to exist in at least three states of differing activity: active, primed and poised enhancers (Calo and Wysocka, 2013), such that only a subset of bound regions (active enhancers) play a role in gene regulation in a given cellular context. Although different states are distinguished by differential patterns of histone modifications and transcriptional regulator recruitment, systematically classifying the state of an enhancer remains a challenge (Zentner et al., 2011; Catarino and Stark, 2018). Given an active enhancer, it is possible to predict tissues in which it is active (Pennacchio et al., 2007; Li et al., 2018). However, these methods do not address the problem of predicting the states of enhancers in a specific tissue.

Modern genomic data is highly heterogeneous and may contain continuous (e.g. histone modification levels), binary (e.g. TF-binding) and discrete features (e.g. counts of methylated sites) and, in principle, supervised machine learning methods can be used to identify active enhancers with integrated heterogeneous genomics data. Enhancer activity, however, is highly tissue-specific (Bulger and Groudine, 2011), and there are few tissues for which large numbers of active enhancers have been identified, limiting the application of recent supervised enhancer prediction approaches such as DECRES (Li et al., 2018) and REPTILE (He et al., 2017) to identify active enhancers in tissues of interest. Furthermore, currently, supervised methods have not been designed to predict enhancer states other than active and inactive enhancers.

Clustering (or unsupervised) techniques could, in principle, identify clusters of genomic regions that are enriched with different states of enhancers without large training sets. Unsupervised methods like ChromHMM (Ernst and Kellis, 2012) and Segway (Hoffman et al., 2012) can predict enhancers; however, these methods were designed for genome segmentation rather than enhancer state prediction over enhancer candidates (like TF-bound regions). Ideally, the small numbers of experimentally validated active enhancers should be used if possible: this motivates the development of semi-supervised approaches that can integrate heterogeneous data.

To integrate high-dimensional genomic data and predict enhancer states over enhancer candidates, we developed a variational Dirichlet Process Heterogeneous Mixture (DPHM or an infinite heterogeneous mixture) model that infers Gaussian, Bernoulli and Poisson distributions over continuous, binary and non-negative discrete features, respectively. To take advantage of small labeled training sets where available, we derive a novel variational inference algorithm for a semi-supervised DPHM model that forces a subset of the data (like experimentally validated enhancers) to cluster together. Our Bayesian model also has the advantages that 1) the number of clusters, or enhancer states, is inferred from the data and 2) the number of hyperparameters does not grow with the number of clusters, simplifying inference for heterogeneous data integration. The DPHM model outperformed Gaussian mixture models in clustering synthetic heterogeneous datasets in unsupervised and semi-supervised settings. The DPHM model can also outperform k-means, in certain settings, even when k-means is given the correct number of clusters.

To illustrate the power of the DPHM model to integrate heterogeneous genomic data, we employed it to predict new enhancers based on heterogeneous features from the Encyclopedia of DNA Elements (ENCODE) Project (ENCODE Project Consortium, 2012). We applied the semi-supervised DPHM model to a dataset of 6,209 genomic regions bound by Nkx2-5, a master regulator in heart development (Tanaka et al., 1999; Schott et al., 1998), in embryonic mouse heart with the constraint that a set of known active enhancers have to cluster together. Through our novel variational inference algorithm, 47 new regions clustered with the experimentally known active enhancers in this tissue. Furthermore, we discovered 5 large classes of genomic regions in the data. Some classes, including the class with the known active enhancers, were significantly (q < 0.05) enriched with various biological processes. Another class, enriched for house-keeping genes, appears to contain active promoters. Moreover, each functional class of enhancers was enriched with at least 60 different TF-binding motifs and some motifs can be utilized to discriminate the classes from each other. Our analysis indicates that the semi-supervised DPHM model is a principled Bayesian method for discovering biologically relevant clusters in heterogeneous genomic data in the semi-supervised learning paradigm.

## Methods

### Data

Dupays et al. (2015) identified genomic regions bound by Nkx2-5 using chromatin immunoprecipitation sequencing (ChIP-seq) from embryonic mouse heart tissue. Genomic coordinates for the binding sites were converted from the GRCm37/mm9 to the GRCm38/mm10 genome build. To incorporate information about the functional conservation of the sites, we identified regions of the human genome (GRCh38/hg38 build) whose DNA sequences are alignable to the sites using LiftOver (Hinrichs et al., 2006). We found that 6209/7246 of the binding sites in the mouse genome had human orthologs. 109 of the regions overlapped with experimentally validated developmental enhancers in mouse heart tissue, identified by the VISTA Enhancer Browser (Visel et al., 2007). Features for the regions were generated from ENCODE datasets (ENCODE Project Consortium, 2012) for mouse and human heart tissues. Supplementary Table 1 lists all of the datasets that we used to extract features. We extracted 81 mouse features and 33 human features. Details about feature extraction techniques are provided in the supplementary.

### Dirichlet Process Heterogeneous Mixtures

The DPHM model takes observations with heterogeneous features and clusters them based on similarities between their features. Continuous, binary and non-negative discrete features are assumed to follow Gaussian, Bernoulli and Poisson distributions, respectively, and are assumed to be mutually independent so the conditional likelihood of an observation is the product of distributions, with cluster-specific parameters, for each feature. In this section, we assume the data has *r*_*g*_, *r*_*b*_ and *r*_*p*_ Gaussian, Bernoulli and Poisson features, respectively.

Figure 1 shows the latent variables of the semi-supervised DPHM model. *μ* and *τ* contain the means and precisions of the Gaussian features, respectively, *p* contains the probability parameters for the Bernoulli features and *λ* contains the average rate parameters for the Poisson features. The subscripts on the variables denote their feature indices and associated clusters. For example, *μ*_*tj*_ represents the mean parameter for the *j-*th Gaussian feature and cluster *t*.

**Fig. 1.**
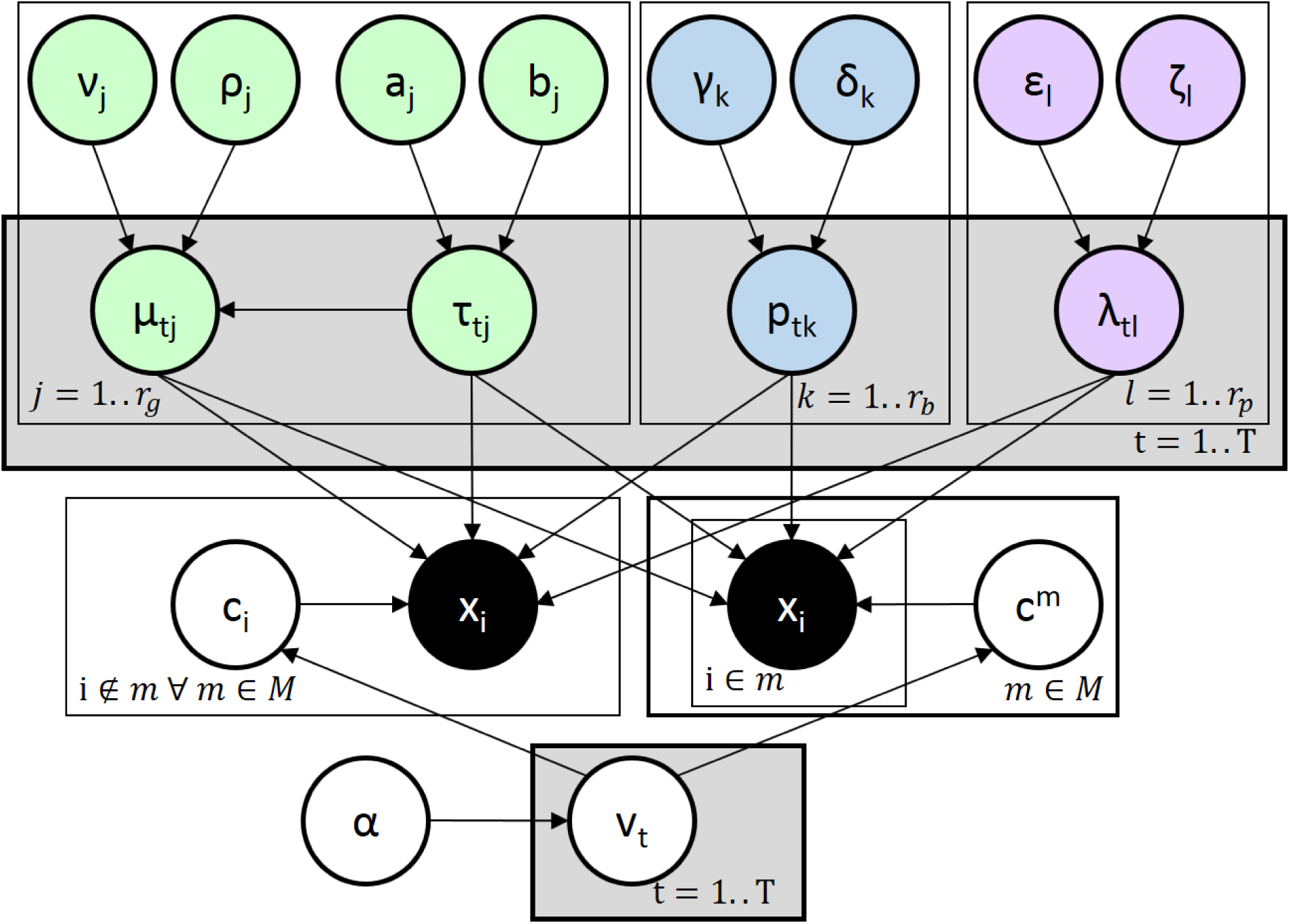
A Bayesian network depicting the dependencies between hyperparameters, latent variables and observations in the DPHM. Each observation, *x*_*i*_, depends on their cluster assignment (*c*_*i*_ for observations that are not in any must-link constraints and *c*^*m*^ for observations in a must-link constraint *m*) and distribution parameters (*μ, τ, p* and *λ*). Distribution parameters depend on parameters of the NGBG prior (*ν*, *ρ*, *a, b*, *γ*, *δ*, *ε* and *ζ*). Distribution parameters and hyperparameters related to Gaussian, Bernoulli and Poisson features are green, blue and purple, respectively. Each cluster assignment depends on the *v*_*t*_s that are generated through the stick-breaking construction. *M* represents a set of must-link constraints for semi-supervised clustering.

#### Priors

Distribution parameters (*μ, τ, p* and *λ*) are drawn from their conjugate priors (Bishop, 2006). The conjugate priors for Gaussian, Bernoulli and Poisson distributions are NormalGamma (NG), Beta and Gamma distributions, respectively. Since features are assumed to be mutually independent, the joint conjugate prior for the distribution parameters can be expressed as:

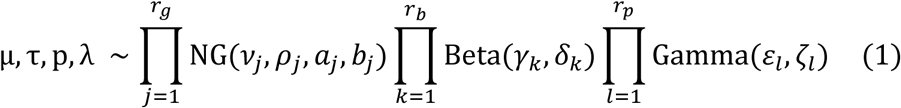

where hyperparameters *ν*_*j*_, *ρ*_*j*_, *a*_*j*_ and *b*_*j*_ control the prior mean, precision, shape and rate parameters, respectively, of the NG distribution that generates the mean and precision for the *j*-th Gaussian feature. *γ*_*k*_ and *δ*_*k*_ represent the prior shape parameters of the Beta distribution that creates the probability parameter for the *k*-th Bernoulli feature. *ε*_*l*_ and *ζ*_*l*_ are the prior shape and rate parameters, respectively, for the Gamma distribution that generates the average rate parameter for the *l*-th Poisson feature. The conjugate prior eases computations in the algorithm because the cluster-specific posterior distributions of *μ, τ, p* and *λ* will be members of the conjugate prior’s family; we refer to this family as ‘NGBG’.

Cluster assignments (denoted by *c*) for each observation are drawn from multinomial distributions, and their prior parameters are mixing weights for the clusters. Mixing weights are constructed through the truncated stick-breaking process (Ishwaran and James, 2001) that sets an upper bound, *T*, on the number of clusters. Mixing weights are completely determined by stick-breaking variables 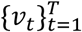 that depend on *α*, a hyperparameter that controls cluster sizes. Although there are *T* clusters, some of them will be empty if *T* is large enough and can be ignored in downstream analyses. Cluster assignments, stick-breaking variables and distribution parameters form the latent variable space, while *α* and parameters of the NGBG prior form the hyperparameter space of the DPHM model.

#### Variational Inference for Semi-supervised DPHM Models

We use variational inference (Beal, 2003; Blei and Jordan, 2006) to fit the DPHM model to a dataset *X*. Variational inference approximates the true posterior of the latent variables with a variational distribution *q* by maximizing the evidence lower bound (ELBO):

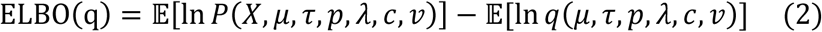

We optimize over the mean-field variational family so *q* factorizes into the product of variational densities for the latent variables. The variational density for a cluster assignment is parameterized by a ‘cluster probability vector’ that contains probabilities of the corresponding observation belonging to the different clusters; each observation is assigned the cluster associated with the maximum probability in its cluster probability vector. We use *φ*_*i*_ to denote the cluster probability vector of the *i*-th observation. The learning task of variational inference is to find variational densities that maximize the ELBO; this can be accomplished through coordinate ascent variational inference (Bishop, 2006). Lim and Wang (2018) derive variational density updates for latent variables of the Dirichlet Process Gaussian Mixture (DPGM) as shown in lines 4, 5 and 8 of Algorithm 1. We use these updates for the variational densities of *μ, τ, c* and *v* in our DPHM model but we must update Bernoulli and Poisson parameters as well. We derived the coordinate-optimal updates for *q*(*p*_*tk*_) and *q*(*λ*_*tl*_) which represent the variational densities of the Bernoulli and Poisson parameters, respectively:

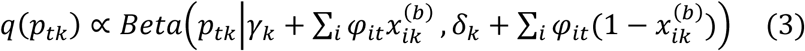

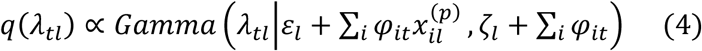

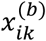 is the *k*-th Bernoulli feature of the *i*-th observation and 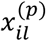 is the *l*-th Poisson feature of the *i*-th observation. *φ*_*it*_ is the probability of the *i*-th observation belonging to cluster *t*. We apply these updates for all *t* = 1..*T*, *k* = 1..*r*_*b*_ and *l* = 1..*r*_*p*_.

We derived a novel variational inference algorithm to fit DPHM models with must-link constraints, outlined in Algorithm 1. In the algorithm, 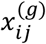 is the *j*-th Gaussian feature of the *i*-th observation and *c*_*-i*_ represents a vector of all cluster assignments except *c*_*i*_. We define a must-link constraint as a set of indices for observations that must cluster together. In Figure 1 and Algorithm 1, *M* denotes a set of must-link constraints because the model can support multiple must-link constraints. While there are variational inference algorithms for semi-supervised classification models (Kingma et al., 2014), with finite numbers of classes, and Gibbs samplers for semi-supervised infinite mixture models (Vlachos et al., 2009), our work presents the first variational inference algorithm designed for semi-supervised infinite mixtures. In our algorithm, all observations in a particular must-link constraint are associated with a single cluster assignment variable, as shown in Figure 1, so they are all assigned the same cluster. In each iteration of coordinate ascent, our algorithm updates variational densities of global latent variables (*μ, τ, p, λ* and *v*) and uses their expectations to calculate cluster probability vectors. For each must-link constraint *m*, with observations *X*^*m*^ = {*x*_*i*_}_*i∈m*_ and cluster assignment *c*^*m*^ (the single cluster assignment variable for all observations in *m*), we derived the coordinate-optimal variational density of *c*^*m*^:

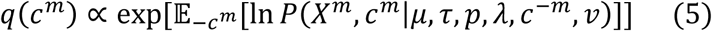

where 𝔼_−*c*^*m*^_ is the expectation with respect to all latent variables except *c*^*m*^ and *c*^−*m*^ is a vector of all cluster assignments except *c*^*m*^. We obtain the cluster probability vector for the observations in *m* by evaluating the variational density over different clusters. Derivations for variational density updates are provided in the supplementary information.

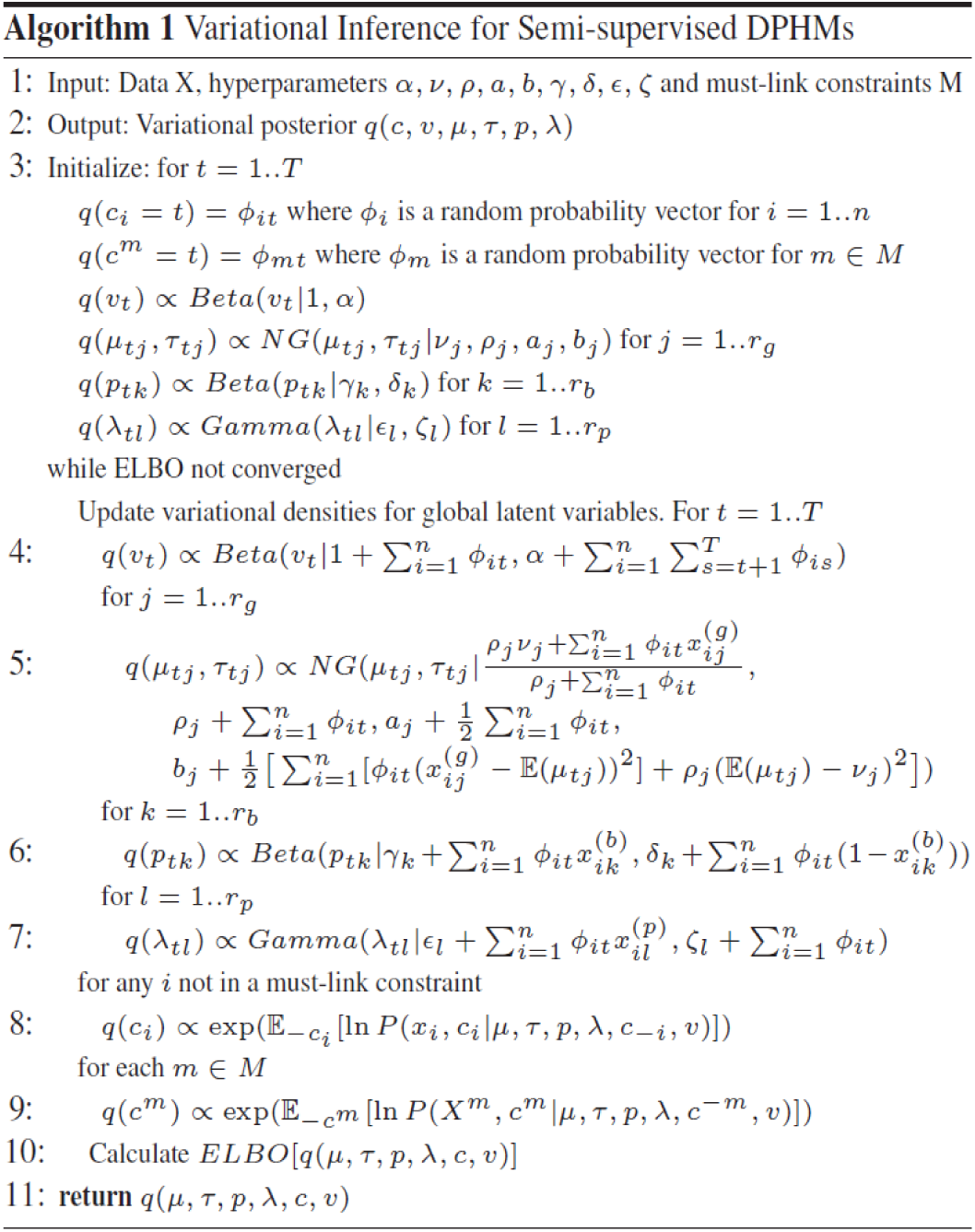

## Results

### Benchmarking the DPHM Against Other Methods

To test whether the DPHM can outperform clustering models that assume Gaussian distributions over all features, we compared it to a finite Gaussian Mixture Model (GMM) and DPGM with the synthetic datasets (supplementary section 2.1.1). The DPHM model achieved significantly (Mann-Whitney U test p < 0.05) higher Adjusted Rand Indices (ARIs) (Hubert and Arabie, 1985) than the GMM and DPGM on all synthetic datasets in unsupervised and semi-supervised settings (Supplementary Figure 1A, p-values shown in Supplementary Table 2), which shows the advantage of incorporating Bernoulli and Poisson distributions into the DPHM model. To determine if the DPHM can outperform a distance-based clustering algorithm, we compared it to k-means with the synthetic datasets and gave k-means the correct numbers of clusters. Against unsupervised k-means, the unsupervised DPHM achieved significantly higher ARIs on the 10 and 25-cluster datasets but there was no significant difference on the 50-cluster dataset. Against constrained k-means (Wagstaff et al., 2001), the semi-supervised DPHM achieved significantly higher ARIs on the 25-cluster dataset but there were no significant differences on the 10 and 50-cluster datasets. The performances of the DPHM and k-means were comparable on certain datasets and settings when k-means was given the correct numbers of clusters.

Since constrained k-means performed well on the synthetic datasets, we applied the DPHM and constrained k-means to a biological dataset (Nkx2-5 dataset) to evaluate their power to predict held-out VISTA enhancers (supplementary section 2.1.2). For each training set, we applied constrained k-means with different values of *k* and initial parameters and picked the setting which maximized the average silhouette (Rousseeuw, 1987). Across all training sets, constrained k-means predicted positive clusters that were an order of magnitude larger than the DPHM (Figure 2A) and achieved higher sensitivity lower bounds (SLBs) than the DPHM, across training set sizes (Figure 2B). The DPHM achieved higher precision lower bounds (PLBs) than constrained k-means (Mann-Whitney U test p < 0.05 for training set sizes between 10 and 90, inclusive, see Supplementary Table 3 for p-values) (Figure 2C). Taken together, these results show that regions appearing in the DPHM’s positive cluster are more likely to be experimentally validated enhancers and that constrained k-means and the DPHM have qualitatively different predictive performances.

**Fig. 2.**
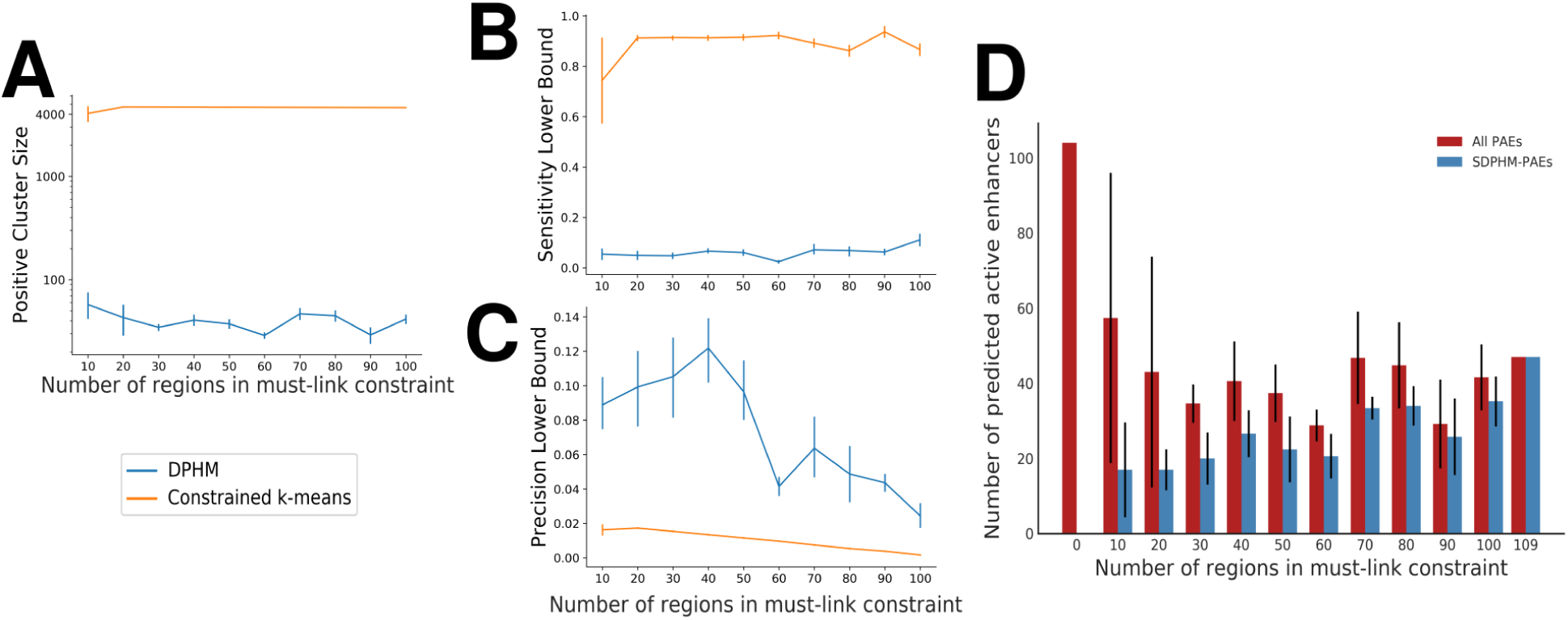
(A) Sizes of positive clusters (the cluster with the training set) predicted by the DPHM and constrained k-means across training set sizes. The y-axis is log-scaled. For each training set size, both models were run 5 times with 5 different training sets. The plot shows mean positive cluster sizes and error bars represent one standard deviation across training sets. (B) Sensitivity lower bounds achieved by the DPHM and constrained k-means with different training set sizes. The plots shows the mean and error bars representing one standard deviation across training sets. (C) Precision lower bounds achieved by the DPHM and constrained k-means with different training set sizes. The plot shows the mean and error bars representing one standard deviation across training sets. (D) A comparison of the numbers of PAEs from the unsupervised DPHM model (with 0 regions in the must-link constraint) and semi-supervised DPHM models with differing training set sizes. For semi-supervised models with 10 to 100 regions in their training set, we randomly sampled their training sets, from the VISTA enhancers, 5 times and calculated the average number of PAEs, across the samples, for each training set size (red bars). In addition, for each training set size, we calculated the average number of PAEs that were also in the set of SDPHM-PAEs (blue bars). Standard deviations, for the numbers of PAEs and SDPHM-PAEs, across sampled training sets were also calculated (black lines).

Finally, we tested the DPHM in a fully supervised learning paradigm. To do so, we considered all VISTA enhancers as the ‘positive set’ and all other regions as the ‘negative set’ and trained standard machine learning classifiers (supplementary section 2.2.3). To fit the parameters of the variational distribution of the fully supervised DPHM model, we created two must-link constraints corresponding to positives and negatives. We then allow all the VISTA enhancers to join either the positive or negative class. We evaluate the sensitivity of the methods in training and held-out VISTA enhancers. In this context, the DPHM is a generative model that assumes independence among features, and as expected, we found that the DPHM achieved similar performance to Naive Bayes (Supplementary Figure 1B). Of the methods we tried, AdaBoost showed the best performance in the fully supervised paradigm, achieving perfect sensitivities (Supplementary Figure 1B).

### Sensitivity to the Training Set

We performed a sensitivity analysis (supplementary section 2.2) to analyze how predicted active enhancers (PAEs) change as we tune the training set of the DPHM. We applied a semi-supervised DPHM model to the Nkx2-5 dataset and trained the model on the complete training set. It discovered 47 PAEs [hereafter referred to as the semi-supervised DPHM’s predicted active enhancers (SDPHM-PAEs)] and most displayed epigenetic marks (Ernst et al., 2011) of active enhancers (Supplementary Figure 2A). Gene ontology (GO) enrichment analysis (supplementary section 2.3) revealed that the nearest genes of the SDPHM-PAEs were significantly enriched for expression in heart ventricle during postnatal development in mouse (q = 3.98 × 10^−2^), suggesting that the semi-supervised model may predict heart-specific active enhancers. One SDPHM-PAE, located 1,322bp upstream from the transcription start site (TSS) of *Actc1* (a gene that transcribes cardiac alpha-actin), was previously functionally validated and confirmed to drive *Actc1* expression (Fleischmann et al., 1998). Sensitivity analysis showed that the DPHM model can predict over half of the SDPHM-PAEs with 70 or more VISTA enhancers in the must-link constraint (Figure 2D). Furthermore, the SDPHM-PAE near *Actc1* was predicted to be an active enhancer for at least 3/5 training set samples with 70 or more VISTA enhancers in the must-link constraint.

For comparison, we applied an unsupervised DPHM to the Nkx2-5 dataset and found that the cluster with the most VISTA enhancers had 17 VISTA enhancers and 104 PAEs. These PAEs had some marks of active enhancers (Supplementary Figure 2B) although their average H3K27 acetylation (K27ac) signal (minmax-scaled K27ac features averaged across time points and PAEs) was significantly (one-sided t-test p = 8.81 × 10^−4^) lower than the average K27ac signal of SDPHM-PAEs. The nearest genes of the PAEs from the unsupervised model were not significantly (q < 0.05) enriched with any GO terms, suggesting that the unsupervised model could not predict enhancers near heart-specific genes. The results for the semi-supervised and unsupervised DPHM models indicate that the training data may allow the DPHM model to predict heart-specific enhancers with stronger activity.

### Clustering the Clusters

The semi-supervised DPHM model, that was trained on the complete training set, found 47 clusters in the Nkx2-5 dataset. In addition to the 47 SDPHM-PAEs, other clusters also contained regions with characteristics of active enhancers. Furthermore, groups of clusters shared characteristics of different enhancer states. We wanted to group similar clusters together using another clustering model. To this end, we represented each cluster (from the semi-supervised DPHM model that was trained on the complete training set) with a feature vector of expected values over its cluster-specific distribution parameters; all the values are continuous. We applied a DPGM (’dphmix’ runs a DPGM when all features are continuous) with α = 1 to group the clusters and found 14 classes (clusters of the original 47 clusters). Since the VISTA enhancers were forced to cluster together, they appeared in the same class. There are five large (over 500 regions) classes that we number from 1 to 5 as shown in Figure 3A. We performed GO enrichment analysis (supplementary section 2.3) on each class to identify possible functions of their regions. Here, we analyze an interesting large class (class 2). Analyses for the other four large classes are provided in the supplementary information. In particular, class 3 contains regions with characteristics of active promoters (Ernst et al., 2011) and class 4 contains regions appearing to be inactive regions or decommissioned enhancers (Pradeepa, 2017).

**Fig. 3.**
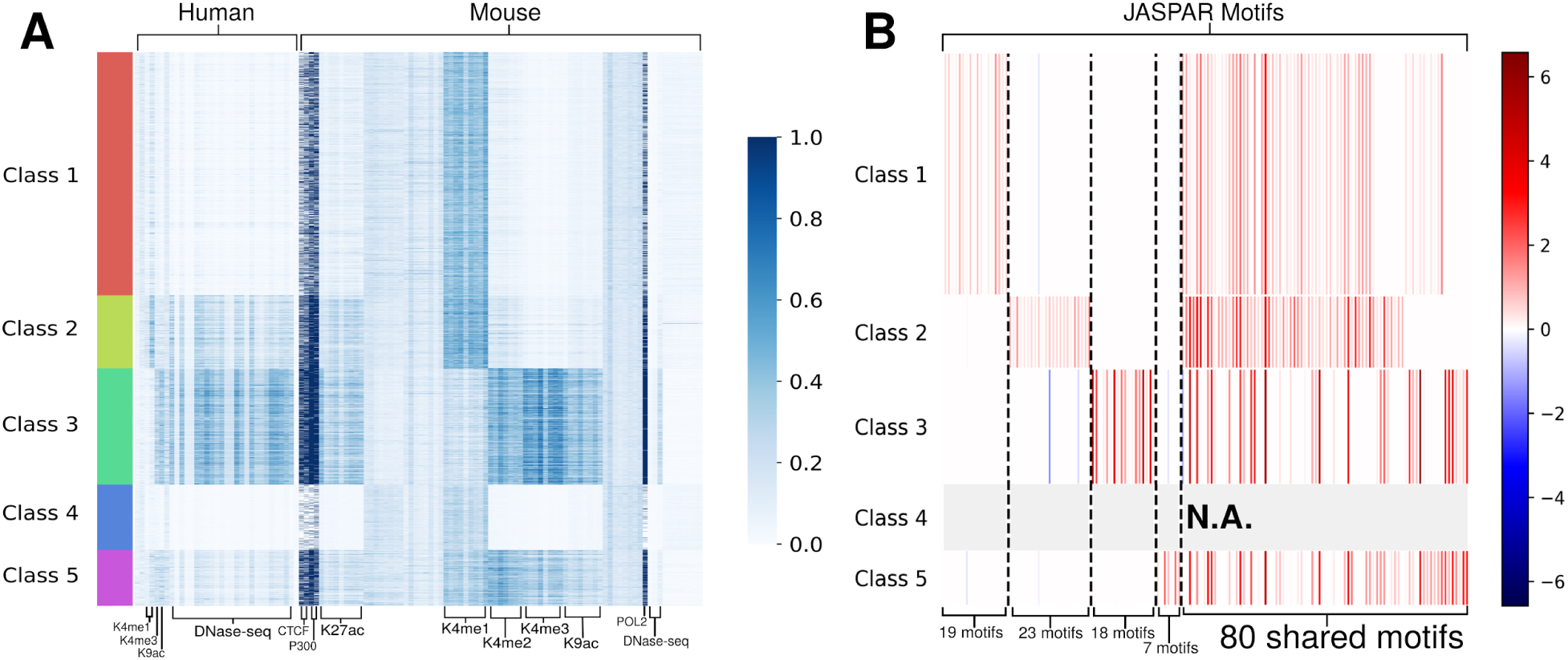
(A) A heatmap showing the regions in the large classes. Each column represents an ENCODE feature in heart tissues and they were minmax-scaled to be between 0 and 1. (B) A heatmap showing the regression coefficients of JASPAR motifs in classes 1, 2, 3 and 5 compared to class 4 (predicted inactive regions). The vertical sizes of each row are proportional to the number of regions in the class corresponding to the rows in panel (A). Only motifs that have a significant (permutation test p < 0.05) and positive coefficient for at least one class compared to class 4 are included. Coefficients for class 4 are not available since it was used as the reference class for all the regressions.

We associated class 2 with active enhancers as it contains the cluster with all of the VISTA enhancers. We refer to this class as the DPHM’s active enhancer class (DPHM-AEC). It also contains 4 other clusters for a total of 741 regions. It has the highest average H3K4 mono-methylation (K4me1) signal (minmax-scaled mouse K4me1 features averaged across time points and regions in the class) and second highest average K27ac signal in embryonic mouse tissue among all classes. Consistent with typical active enhancers, P300 and POL2 were bound to the vast majority of the regions in adult tissue (Spicuglia and Vanhille, 2012). GO annotation analysis on genes near (within 1Mb) regions from the DPHM-AEC revealed 100 significantly enriched biological processes. To ensure that the observed enrichments were not biased by VISTA enhancers, we conducted GO enrichment analysis on the regions in the class that are not in VISTA. The regions were still significantly enriched with 6/100 of the aforementioned processes, suggesting that they may be novel active enhancers with similar functions as VISTA enhancers.

#### Comparison to Classes Discovered by Constrained K-means

To determine if constrained k-means can discover classes of enhancers, we applied constrained k-means on the Nkx2-5 dataset with a must-link constraint on all 109 VISTA enhancers. Maximizing the average silhouette yielded *k*=2 classes (Supplementary Figure 3A). One class contains regions that appear to be active promoters while the other class contains all other regions, including the VISTA enhancers (Supplementary Figure 3B). Neither class was significantly (q < 0.05) enriched with any biological processes. Hence, if k is chosen to maximize the average silhouette, constrained k-means may separate enhancers from promoters but it could not discover different states of enhancers. This illustrates the advantage of the DPHM, which appears to identify multiple biologically relevant classes with characteristics of different enhancer states.

Next, we applied constrained k-means with *k*=14 (to match the number of classes found by the DPHM). Constrained k-means found 3 large classes. Two of these contain a mixture of regions with characteristics of poised, primed, weak, inactive and decommissioned enhancers (Supplementary Figure 3C, red and cyan bars). The other large class had the VISTA enhancers so we refer to this class as the constrained k-means’ active enhancer class (CKM-AEC). Regions in the CKM-AEC had characteristics of active enhancers (Supplementary Figure 3C). The genes near regions from the CKM-AEC were not significantly (q < 0.05) enriched with any processes that were enriched in VISTA enhancers whereas 2 heart-specific processes enriched in the DPHM-AEC were enriched in VISTA enhancers (Supplementary files: class2_noVISTA_SigProcesses and ckm_aec_noVISTA_SigProcesses), supporting the idea that the DPHM has more power to predict heart-specific active enhancers even when k was given. Most of the regions that appeared to be active promoters (class 3 found by the DPHM) are now dispersed over 5 other classes found by constrained k-means. The analysis reveals that constrained k-means can identify different classes of enhancers, given the right k, but the classes appear less specific than those found by the DPHM.

#### Motif Enrichment Analysis of Classes

We performed motif enrichment analysis (supplementary section 2.4) to identify differentially enriched motifs, which were conserved across 6 mammals, between classes found by the DPHM. Each class was enriched with at least 60 different conserved motifs, compared to class 4 (predicted inactive regions). In particular, 86 motifs were enriched in the DPHM-AEC and their TFs were significantly (q < 0.05) enriched for 17 biological processes, including regulation of muscle system process (q = 6.59 × 10^−3^). The motifs for other classes were not significantly enriched with any processes, suggesting that the TFs associated with the 23 motifs (or a subset of them) uniquely enriched in the DPHM-AEC may preferentially bind to active enhancers and facilitate biological processes in the developing mouse heart. While 80 motifs are enriched in multiple classes (shared motifs), each class is uniquely enriched with at least 7 motifs (Figure 3B) suggesting that many different TFs may be required to determine enhancer states. We repeated this analysis using multinomial regression and found similar results (supplementary figure 4).

## Discussion

We derived the semi-supervised variational DPHM model to cluster heterogeneous data with must-link constraints using a Bayesian framework. We showed that the DPHM and constrained k-means have qualitatively different predictive performances on biological data and that the DPHM achieves higher precision lower bounds, which is important in enhancer prediction because false positives are costly in experimental validation. We used the DPHM to cluster genomic regions bound by a master regulator in embryonic mouse heart based on heterogeneous ENCODE features and predicted the most similar regions to a set of known active enhancers. Further studies will determine whether other epigenomics datasets (Roadmap Epigenomics Consortium et al., 2015; Noguchi et al., 2017) and positive enhancers identified through self-transcribing active regulatory region sequencing (STARR-seq) (Liu et al., 2017) can potentially be utilized to train DPHMs and predict states of new enhancer candidates.

Clustering the clusters revealed multiple functional classes of enhancers in the Nkx2-5 ChIP-seq dataset, and motif enrichment analysis suggested that many different motifs may be required to discriminate the classes from each other. Experimental validation is required to determine if the motifs uniquely enriched in specific classes control different enhancer states and whether multiple binding events are required for enhancers to transition from an inactive state to a functional state. However, there are examples of multiple TF binding events driving expression of the Hbb gene to facilitate cell fate transitions (Capellera-Garcia et al., 2016; Mitchell et al., 2012).

Our semi-supervised DPHM approach solves a different problem than other enhancer prediction methods (Ernst and Kellis, 2012; Hoffman et al., 2012; He et al., 2017; Li et al., 2018) as it predicts enhancer states from a set of bound regions. In the fully supervised paradigm, we found AdaBoost was better at predicting held-out VISTA enhancers, but this is not the recommended use case for our model: the DPHM was designed to predict an unbounded number of enhancer states. Supervised enhancer classifiers can only predict classes that are present in the training set. To separate active enhancers from inactive enhancers, they require labelled training examples of active and inactive enhancers. While training regions can be labelled based on the presence of certain histone marks, TF binding or enhancer RNA expression, these are neither necessary nor sufficient for enhancer activity (Catarino and Stark, 2018). The advantage of the semi-supervised approach is to predict multiple classes even when the training set only includes validated active enhancers. Consistent with this, we also showed that constrained k-means can predict enhancer classes, but requires a priori information about the number of classes. Moreover, although the PAEs from the unsupervised DPHM model were weaker candidates for active enhancers compared to the SDPHM-PAEs, the unsupervised model discovered PAEs with some active enhancer marks so it can still be used for enhancer clustering.

One of the advantages of an infinite mixture model over a finite mixture model is that the number of hyperparameters in an infinite mixture model only scales with the number of features, whereas, in a finite mixture model, it scales with the number of features and clusters. Our DPHM model has 1 + 4*r*_*g*_ + 2*r*_*b*_ + 2*r*_*p*_ hyperparameters, while a GMM, with a diagonal covariance matrix, would have *k* − 1 + 2*k*(*r*_*g*_ + *r*_*b*_ + *r*_*p*_), where *k* is the number of clusters. Hence, the DPHM model reduces the number of hyperparameters by a factor of order *k*, which can be substantial if *k* is large. Instead of variational inference, DPHMs can also be fit with Markov Chain Monte Carlo methods (Neal, 2000), however, convergence is difficult to assess and not guaranteed in a finite number of iterations. We implemented a Gibbs sampler (Vlachos et al., 2009) to fit the semi-supervised DPHM but found that our implementation was numerically unstable when the data had a large number of features.

Further work is required to determine whether our DPHM model is viable for clustering regions genome-wide based on chromatin state. The algorithm is *O*(Tn) as it passes through the entire dataset and all clusters to update cluster probability vectors, but would still require a large amount of time for large datasets as it has to complete a full pass over the dataset at each iteration of coordinate ascent. Moreover, there is substantial overhead in gathering files, extracting features for every genomic region and exhaustively testing different priors. Stochastic variational inference could speed up the algorithm for larger datasets as it allows some variational parameters to be updated based on subsamples of the data (Hoffman et al., 2013).

The purpose of clustering the clusters was to reduce the number of clusters, but, in principle, this can also be accomplished by tuning α. For large datasets, however, α has a marginal effect on the number of clusters, because variational densities for most of the stick-breaking variables are primarily determined by cluster probabilities rather than α (line 4 of Algorithm 1). The clustering of clusters is a novel, model-based method to hierarchically cluster data in a multi-way tree structure. Blundell *et al.* (2010) derived a hierarchical clustering algorithm that produces multi-way trees, but the algorithm is *O*(n^2^ log n), so it may have difficulties scaling to large data. In contrast, even for hierarchical clustering, our method is still *O*(Tn), as each layer of clustering is done independently and only adds terms of order *n* to the time complexity. Nonetheless, our method is heuristic and further work is needed to create a principled, scalable Bayesian hierarchical clustering model.

## Conclusion

In this paper, we derived the DPHM model to cluster genomic regions bound by a master regulator in embryonic mouse heart based on heterogeneous features. In addition, we derived a variational inference algorithm to force known active enhancers to cluster together. The semi-supervised DPHM model discovered 47 regions that were similar to the known active enhancers and near heart-specific genes. Furthermore, we clustered the clusters from the DPHM model to find 5 large classes of enhancers with distinct patterns over their features. The class with the known active enhancers was enriched with heart-specific biological processes and TF-binding motifs that are important for muscle system processes. The other functional classes of enhancers were also enriched with many different TF-binding motifs. Our results show that the semi-supervised DPHM model provides a principled Bayesian method for clustering heterogeneous data with small training sets.

## Supporting information

class2_noVISTA_SigProcesses

ckm_aec_noVISTA_SigProcesses

## Acknowledgements

We thank Rachel Chan, Lee Zamparo, Michael Hoffman and the members of the Moses Lab for helpful comments on the manuscript.

## References

Beal, M. (2003). Variational Algorithms for Approximate Bayesian Inference. Ph.D. thesis, Gatsby Computational Neuroscience Unit, University College London.

Bishop, C. M. (2006). Pattern Recognition and Machine Learning. Springer New York. Blei, D. M. and Jordan, M. I. (2006). Variational inference for dirichlet process mixtures. Bayesian analysis, 1(1), 121–143.

Blundell, C. et al. (2010). Bayesian rose trees. In Proceedings of the Twenty-Sixth Conference on Uncertainty in Artificial Intelligence, UAI’10, pages 65–72, Arlington, Virginia, United States. AUAI Press.

Bulger,M. and Groudine,M. (2011). Functional and mechanistic diversity of distal transcription enhancers. Cell, 144(3), 327–339.

Calo, E. and Wysocka, J. (2013). Modification of enhancer chromatin: What, how, and why? Molecular Cell, 49(5), 825–837.

Capellera-Garcia, S. et al. (2016). Defining the minimal factors required for erythropoiesis through direct lineage conversion. Cell Rep., 15(11), 2550–62.

Catarino, R. R. and Stark, A. (2018). Assessing sufficiency and necessity of enhancer activities for gene expression and the mechanisms of transcription activation. Genes & Development, 32(3-4), 202–223.

Dupays, L. et al. (2015). Sequential binding of meis1 and nkx2-5 on the popdc2 gene: A mechanism for spatiotemporal regulation of enhancers during cardiogenesis. Cell Reports, 13(1), 183–195.

ENCODE Project Consortium (2012). An integrated encyclopedia of dna elements in the human genome. Nature, 489(7414), 57–74.

Ernst, J. and Kellis, M. (2012). Chromhmm: automating chromatin-state discovery and characterization. Nature methods, 9(3), 215–6.

Ernst, J. et al. (2011). Mapping and analysis of chromatin state dynamics in nine human cell types. Nature, 473(7345), 43–49.

Fleischmann, M. et al. (1998). Cardiac specific expression of the green fluorescent protein during early murine embryonic development. FEBS letters, 440(3), 370–6.

He, Y. et al. (2017). Improved regulatory element prediction based on tissue-specific local epigenomic signatures. Proceedings of the National Academy of Sciences, 114(9), E1633–E1640.

Hinrichs, A. et al. (2006). The ucsc genome browser database: update 2006. Nucleic acids research, 34(suppl 1), D590–D598.

Hoffman, M. D. et al. (2013). Stochastic variational inference. The Journal of Machine Learning Research, 14(1), 1303–1347.

Hoffman, M. M. et al. (2012). Unsupervised pattern discovery in human chromatin structure through genomic segmentation. Nature methods, 9(5), 473–476.

Hubert, L. and Arabie, P. (1985). Comparing partitions. Journal of Classification, 2(1), 193–218.

Ishwaran, H. and James, L. F. (2001). Gibbs sampling methods for stick breaking priors. Journal of the American Statistical Association, 96, 161–173.

Kingma, D. P. et al. (2014). Semi-supervised learning with deep generative models. In Advances in Neural Information Processing Systems, pages 3581–3589.

Li, Y. et al. (2018). Genome-wide prediction of cis-regulatory regions using supervised deep learning methods. BMC Bioinformatics, 19(202).

Lim, K.-L. and Wang, H. (2018). Fast approximation of variational bayes dirichlet process mixture using the maximization–maximization algorithm. International Journal of Approximate Reasoning, 93, 153–177.

Liu, Y. et al. (2017). Functional assessment of human enhancer activities using whole-genome STARR-sequencing. Genome Biology, 18(1), 219.

Mitchell, J. A. et al. (2012). Nuclear rna sequencing of the mouse erythroid cell transcriptome. PLoS One, 7(11), e49274.

Neal, R.M. (2000). Markov chain sampling methods for dirichlet process mixture models. Journal of computational and graphical statistics, 9(2), 249–265.

Noguchi, S. et al. (2017). Fantom5 cage profiles of human and mouse samples. Sci Data., 4(170112).

Pennacchio, L. A. et al. (2007). Predicting tissue-specific enhancers in the human genome. Genome Research, 17(2), 201–211.

Pradeepa, M. M. (2017). Causal role of histone acetylations in enhancer function. Transcription, 8(1), 40–47.

Roadmap Epigenomics Consortium et al. (2015). Integrative analysis of 111 reference human epigenomes. Nature, 518(7539), 317–330.

Rousseeuw, P. J. (1987). Silhouettes: A graphical aid to the interpretation and validation of cluster analysis. Journal of Computational and Applied Mathematics, 20, 53–65.

Schott, J.-J. et al. (1998). Congenital Heart Disease Caused by Mutations in the Transcription Factor NKX2-5. Science, 281(5373), 108–111.

Spicuglia, S. and Vanhille, L. (2012). Chromatin signatures of active enhancers. Nucleus, 3(2), 126–131.

Tanaka, M. et al. (1999). The cardiac homeobox gene Csx/Nkx2.5 lies genetically upstream of multiple genes essential for heart development. Development, 126(6), 1269–1280.

Visel, A. et al. (2007). Vista enhancer browser–a database of tissue-specific human enhancers. Nucleic acids research, 35(Database issue), D88–92.

Vlachos, A. et al. (2009). Unsupervised and constrained dirichlet process mixture models for verb clustering. In Proceedings of the Workshop on Geometrical Models of Natural Language Semantics, GEMS ’09, pages 74–82, Stroudsburg, PA, USA. Association for Computational Linguistics.

Wagstaff, K. et al. (2001). Constrained k-means clustering with background knowledge. In Proceedings of the Eighteenth International Conference on Machine Learning, ICML ’01, pages 577–584, San Francisco, CA, USA. Morgan Kaufmann Publishers Inc.

Zentner, G. E. et al. (2011). Epigenetic signatures distinguish multiple classes of enhancers with distinct cellular functions. Genome Res., 21, 1273–1283.

