## Supplementary Information for "Variational Infinite Heterogeneous Mixture Model for Semi-supervised Clustering of Heart Enhancers"

#### Contents

**1 Derivations for Variational Density Updates** 1

**2 Methods** 3
- 2.1 Feature Extraction 3
- 2.2 Benchmarking 4
  - 2.2.1 Comparison to unsupervised and semi-supervised methods on synthetic data 4
  - 2.2.2 Comparison to constrained k-means on biological data 5
  - 2.2.3 Comparison to supervised classifiers on biological data 5
- 2.3 Sensitivity Analysis 5
- 2.4 GO Enrichment Analysis 5
- 2.5 Motif Enrichment Analysis 6

**3 Results** 6
- 3.1 Classes Found by the DPHM 6
- 3.2 Multinomial Lasso Logistic Regression 7

**4 Reproducing our Clusters** 7

**5 References** 8

**6 Supplementary Tables & Figures** 9

### 1 Derivations for Variational Density Updates

We derived variational density updates for Bernoulli and Poisson parameters by taking partial derivatives of the evidence lower bound (ELBO) with respect to the density of interest. We use $\mu, \tau, p, \lambda, c$ and $v$ to denote parameter vectors. For example, $p$ contains Bernoulli parameters for all features and clusters. Let $M$ represent the set of must-link constraints and $\overline{M}$ be the set of indices of observations that are not in must-link constraints. To derive the coordinate-optimal update for $q(p_{tk})$, the variational density of the Bernoulli parameter for the $k^{th}$ binary feature and cluster t, we write the ELBO in terms of $p_{tk}$.

$$\begin{aligned} ELBO &= \mathbb{E}\big[\ln P(X,\mu,\tau,p,\lambda,c,v)\big] - \mathbb{E}\big[\ln q(\mu,\tau,p,\lambda,c,v)\big] \\ ELBO &= \mathbb{E}\big[\ln P(X,\mu,\tau,p_{-tk},\lambda,c,v)P(p_{tk}|X,\mu,\tau,p_{-tk},\lambda,c,v)\big] \\ &\quad - \mathbb{E}\big[\ln \prod_{i\in\overline{M}} q(c_i) \prod_{m\in M} q(c^m) \prod_{z=1}^{T}[q(v_z)\prod_{j=1}^{r_g} q(\mu_{zj},\tau_{zj})\prod_{y=1}^{r_b} q(p_{zy})\prod_{l=1}^{r_p} q(\lambda_{zl})]\big] \end{aligned} \tag{1}$$

where $p_{-tk}$ is a vector of all Bernoulli parameters except $p_{tk}$.

$$\begin{aligned} ELBO &= \mathbb{E}\big[\ln P(X,\mu,\tau,p_{-tk},\lambda,c,v)\big] + \mathbb{E}\big[\ln P(p_{tk}|X,\mu,\tau,p_{-tk},\lambda,c,v)\big] \\ &\quad - \mathbb{E}\big[\ln q(\mu,\tau,p_{-tk},\lambda,c,v)\big] - \mathbb{E}_{p_{tk}}\big[\ln q(p_{tk})\big] \end{aligned} \tag{2}$$

where $\mathbb{E}_{p_{tk}}$ is the expectation with respect to $p_{tk}$. Since we will take the derivative with respect to $q(p_{tk})$, any term that does not contain $p_{tk}$ can be treated as a constant.

$$ELBO = \mathbb{E}_{p_{tk}}\Big[\mathbb{E}_{-p_{tk}}\big[\ln P(p_{tk}|X,\mu,\tau,p_{-tk},\lambda,c,v)\big]\Big] - \mathbb{E}_{p_{tk}}\big[\ln q(p_{tk})\big] + const. \tag{3}$$

where $\mathbb{E}_{-p_{tk}}$ is the expectation with respect to all latent variables except $p_{tk}$. Evaluating $\mathbb{E}_{p_{tk}}$ for both terms results in the following equation:

$$ELBO = \int q(p_{tk})\mathbb{E}_{-p_{tk}}\big[\ln P(p_{tk}|X,\mu,\tau,p_{-tk},\lambda,c,v)\big]dp_{tk} - \int q(p_{tk})\ln q(p_{tk})dp_{tk} + const. \tag{4}$$

Now, we take the partial derivative with respect to $q(p_{tk})$

$$\frac{\partial ELBO}{\partial q(p_{tk})} = \mathbb{E}_{-p_{tk}}\big[\ln P(p_{tk}|X,\mu,\tau,p_{-tk},\lambda,c,v)\big] - \ln q(p_{tk}) - 1 + const. \tag{5}$$

Additional constants arise from Lagrange multipliers. Setting the partial derivative to 0 and solving for $q(p_{tk})$ leads to:

$$\begin{aligned} q(p_{tk}) &= \exp\Big[\mathbb{E}_{-p_{tk}}\big[\ln P(p_{tk}|X,\mu,\tau,p_{-tk},\lambda,c,v)\big] + const.\Big] \\ q(p_{tk}) &\propto \exp\Big[\mathbb{E}_{-p_{tk}}\big[\ln P(p_{tk}|X,\mu,\tau,p_{-tk},\lambda,c,v)\big]\Big] \end{aligned} \tag{6}$$

Applying Bayes' rule to $P(p_{tk}|X,\mu,\tau,p_{-tk},\lambda,c,v)$ reveals:

$$\begin{aligned} q(p_{tk}) &\propto \exp\big[\mathbb{E}_{-p_{tk}}[\ln P(X,\mu,\tau,p,\lambda,c,v)]\big] \\ &\propto \exp\big[\mathbb{E}_{-p_{tk}}[\ln P(p_{tk}) + \ln P(X|\mu,\tau,p,\lambda,c,v)]\big] \\ &\propto \exp\big[\ln P(p_{tk}) + \mathbb{E}_{-p_{tk}}[\ln \prod_{i\in\overline{M}} P(x_i|\mu,\tau,p,\lambda,c,v) \prod_{m\in M}\prod_{i\in m} P(x_i|\mu,\tau,p,\lambda,c,v)]\big] \end{aligned} \tag{7}$$

The likelihood of $x_i$ is expressed as:

$$P(x_i|\mu,\tau,p,\lambda,c,v) = \prod_{z=1}^{T}\Bigg[\Big[\prod_{j=1}^{r_g} N(x_{ij}^{(g)}|\mu_{zj},\tau_{zj}) \prod_{y=1}^{r_b} Bern(x_{iy}^{(b)}|p_{zy}) \prod_{l=1}^{r_p} Poi(x_{il}^{(p)}|\lambda_{zl})\Big]^{I(c_i=z)}\Bigg] \tag{8}$$

If $i \in m$ for some must-link constraint $m$ then we replace $c_i$ with $c^m$. Substituting the log likelihoods and ignoring terms that do not depend on $p_{tk}$ yields:

$$\begin{aligned} q(p_{tk}) &\propto \exp\big[\ln P(p_{tk}) + \sum_{i\in\overline{M}} \mathbb{E}_{-p_{tk}}[I(c_i=t)]\ln P(x_{ik}^{(b)}|p_{tk}) + \sum_{m\in M}\sum_{i\in m}\mathbb{E}_{-p_{tk}}[I(c^m=t)]\ln P(x_{ik}^{(b)}|p_{tk})]\big] \\ &\propto \exp\big[\ln P(p_{tk}) + \sum_{i\in\overline{M}} \phi_{it}\ln P(x_{ik}^{(b)}|p_{tk}) + \sum_{m\in M}\sum_{i\in m}\phi_{mt}\ln P(x_{ik}^{(b)}|p_{tk})]\big] \end{aligned} \tag{9}$$

$\phi_{it}$ is the probability of the $i^{th}$ observation belonging to cluster t and $\phi_{mt}$ is the probability of an observation in $m$ belonging to cluster t. This means $\phi_{mt} = \phi_{it}$ for $i \in m$. This allows the equation to simplify:

$$\begin{aligned} q(p_{tk}) &\propto \exp\big[\ln P(p_{tk}) + \sum_i \phi_{it}\ln P(x_{ik}^{(b)}|p_{tk})\big] \\ &\propto \exp\big[(\gamma_k - 1)\ln p_{tk} + (\delta_k - 1)\ln(1-p_{tk}) + \sum_i \phi_{it}[x_{ik}^{(b)}\ln p_{tk} + (1-x_{ik}^{(b)})\ln(1-p_{tk})]\big] \\ &\propto \exp\big[(\gamma_k + \sum_i \phi_{it}x_{ik}^{(b)} - 1)\ln p_{tk} + (\delta_k + \sum_i \phi_{it}(1-x_{ik}^{(b)}) - 1)\ln(1-p_{tk})\big] \\ &\propto Beta\Big(p_{tk}|\gamma_k + \sum_i \phi_{it}x_{ik}^{(b)},\ \delta_k + \sum_i \phi_{it}(1-x_{ik}^{(b)})\Big) \end{aligned} \tag{10}$$

We can apply a similar procedure to derive the update for $q(\lambda_{tl})$, the variational density of the Poisson parameter for the $l^{th}$ non-negative discrete feature and cluster t, to obtain:

$$\begin{aligned} q(\lambda_{tl}) &\propto \exp\big[\mathbb{E}_{-\lambda_{tl}}[\ln P(X,\mu,\tau,p,\lambda,c,v)]\big] \\ &\propto \exp\big[\ln P(\lambda_{tl}) + \sum_i \phi_{it}\ln P(x_{il}^{(p)}|\lambda_{tl})\big] \\ &\propto \exp\big[(\epsilon_l - 1)\ln\lambda_{tl} - \zeta_l\lambda_{tl} + \sum_i \phi_{it}[x_{il}^{(p)}\ln\lambda_{tl} - \lambda_{tl}]\big] \\ &\propto \exp\big[(\epsilon_l + \sum_i \phi_{it}x_{il}^{(p)} - 1)\ln\lambda_{tl} - (\zeta_l + \sum_i \phi_{it})\lambda_{tl}\big] \\ &\propto Gamma\Big(\lambda_{tl}|\epsilon_l + \sum_i \phi_{it}x_{il}^{(p)},\ \zeta_l + \sum_i \phi_{it}\Big) \end{aligned} \tag{11}$$

To derive the coordinate-optimal update for $q(c^m)$ (the variational density of a cluster assignment for a must-link constraint $m$), we take the partial derivatives of the ELBO with respect to $q(c^m = t) = \phi_{mt}$ for all $t = 1..T$.

$$\begin{aligned} ELBO &= \mathbb{E}\big[\ln P(X, \mu, \tau, p, \lambda, c^{-m}, v) P(c^m | X, \mu, \tau, p, \lambda, c^{-m}, v)\big] \\ &- \mathbb{E}\big[\ln \prod_{i \in \overline{M}} q(c_i) \prod_{w \in M} q(c^w) \prod_{z=1}^{T} [q(v_z) \prod_{j=1}^{r_g} q(\mu_{zj}, \tau_{zj}) \prod_{k=1}^{r_b} q(p_{zk}) \prod_{l=1}^{r_p} q(\lambda_{zl})]\big] \end{aligned} \tag{12}$$

$c^{-m}$ is a vector of all cluster assignments except $c^m$.

$$\begin{aligned} ELBO &= \mathbb{E}_{c^m}\Big[\mathbb{E}_{-c^m}\big[\ln P(c^m | X, \mu, \tau, p, \lambda, c^{-m}, v)\big]\Big] - \mathbb{E}_{c^m}\big[\ln q(c^m)\big] + const. \\ ELBO &= \sum_{z=1}^{T} q(c^m = z) \mathbb{E}_{-c^m}\big[\ln P(c^m = z | X, \mu, \tau, p, \lambda, c^{-m}, v)\big] - \sum_{z=1}^{T} q(c^m = z) \ln q(c^m = z) + const. \\ ELBO &= \sum_{z=1}^{T} \phi_{mz} \mathbb{E}_{-c^m}\big[\ln P(c^m = z | X, \mu, \tau, p, \lambda, c^{-m}, v)\big] - \sum_{z=1}^{T} \phi_{mz} \ln \phi_{mz} + const. \\ \frac{\partial ELBO}{\partial \phi_{mt}} &= \mathbb{E}_{-c^m}\big[\ln P(c^m = t | X, \mu, \tau, p, \lambda, c^{-m}, v)\big] - \ln \phi_{mt} - 1 + const. \end{aligned} \tag{13}$$

Setting this to 0 and solving for $\phi_{mt}$ yields:

$$\begin{aligned} \phi_{mt} &\propto \exp\Big[\mathbb{E}_{-c^m}\big[\ln P(c^m = t | X, \mu, \tau, p, \lambda, c^{-m}, v)\big]\Big] \\ &\propto \exp\Big[\mathbb{E}_{-c^m}\big[\ln P(X, \mu, \tau, p, \lambda, c^{-m}, v, c^m = t)\big]\Big] \\ &\propto \exp\Big[\mathbb{E}_{-c^m}\big[\ln P(c^m = t | v) + \ln P(X | \mu, \tau, p, \lambda, c^{-m}, v, c^m = t)\big]\Big] \\ &\propto \exp\Big[\mathbb{E}_{-c^m}\big[\ln P(c^m = t | v) \\ &+ \ln \prod_{i \in \overline{M}} P(x_i | \mu, \tau, p, \lambda, c^{-m}, v, c^m = t) \prod_{w \in M} \prod_{i \in w} P(x_i | \mu, \tau, p, \lambda, c^{-m}, v, c^m = t)\big]\Big] \end{aligned} \tag{14}$$

For the log likelihood, we only need to consider terms that depend on $c^m$ (observations in $m$).

$$\begin{aligned} \phi_{mt} &\propto \exp\Big[\mathbb{E}_{-c^m}\big[\ln P(c^m = t | v) + \ln \prod_{i \in m} P(x_i | \mu, \tau, p, \lambda, c^{-m}, v, c^m = t)\big]\Big] \\ &\propto \exp\Big[\mathbb{E}_{-c^m}\big[\ln P(X^m, c^m = t | \mu, \tau, p, \lambda, c^{-m}, v)\big]\Big] \end{aligned} \tag{15}$$

where $X^m = \{x_i\}_{i \in m}$. In general, $q(c^m) \propto \exp\Big[\mathbb{E}_{-c^m}\big[\ln P(X^m, c^m | \mu, \tau, p, \lambda, c^{-m}, v)\big]\Big]$

### 2 Methods

#### 2.1 Feature Extraction

The mouse features were comprised of 68 continuous features, 5 binary features and 8 ordinal features. The continuous features were composed of 65 histone modification features and 3 DNase-seq features. For histone modification features, we used 65 different chromatin immunoprecipitation sequencing (ChIP-seq) datasets for different histone modifications and time points. For DNase-seq features, we used 3 different DNase-seq datasets for 3 different time points. The binary features were created from 5 different TF ChIP-seq datasets (2 datasets for CTCF binding at 2 different time points, 2 datasets for P300 at 2 different time points and 1 dataset for POL2 binding). Finally, ordinal features were generated from whole-genome bisulfite sequencing (WGBS) data, which measures DNA methylation (DNAm) levels of CpG sites. We used 8 WGBS datasets for 8 different time points. For each observation, continuous features were extracted by taking the maximum histone modification ChIP-seq or DNase-seq signal across its corresponding genomic region in mouse. Binary features were extracted, for each observation, to indicate whether the TF was bound to the corresponding region in mouse (1 if there is a ChIP-seq peak in the region, 0 otherwise). Finally, for each observation, ordinal features were extracted by counting the number of highly methylated (DNAm level $> 0.5$) CpG sites across the corresponding region in mouse.

Human features consisted of 32 continuous features and 1 ordinal feature. Continuous features were composed of 7 histone modification features and 25 DNase-seq features. For histone modification features, we used 7 different ChIP-seq datasets for different histone modifications, time points and samples. For DNase-seq features, we used 25 DNase-seq datasets for different time points and samples. The ordinal feature was created using WGBS data from an embryonic time point. We extracted human features using the same methods that we employed to extract mouse features.

#### 2.2 Benchmarking

##### 2.2.1 Comparison to unsupervised and semi-supervised methods on synthetic data

To show the advantage of the Dirichlet Process Heterogeneous Mixture (DPHM) model in clustering heterogeneous data in unsupervised and semi-supervised settings, we benchmarked its performance against comparators [Dirichlet Process Gaussian Mixtures (DPGMs), finite Gaussian Mixture Models (GMMs) and k-means] on three synthetic datasets with 10, 25 and 50 clusters, respectively. Each dataset had 1000 observations with 10 Gaussian, 10 Bernoulli and 10 Poisson features. For each type of feature, there are 5 informative features that have cluster-specific distributions, and 5 nuisance features that have the same distribution for all clusters. Mean parameters for the Gaussian features were uniformly generated from (-100, 100) and standard deviations were uniformly generated from (0, 10). Probability parameters for Bernoulli features were uniformly generated from (0,1). Average rate parameters for Poisson features were generated uniformly from (0, 100). Performances were evaluated using Adjusted Rand Indices (ARIs) (Hubert and Arabie, 1985).

We benchmarked the DPHM model against finite GMMs and DPGMs with diagonal covariance matrices, and k-means on the three synthetic datasets in an unsupervised setting. As the predictions of each model vary due to parameter initializations, we applied each model 10 times on each dataset with different initializations (10 different initial cluster probability vectors for the DPHMs and DPGMs, 10 different initial means and covariance matrices for the GMMs and 10 different initial centroids for k-means). The GMM and k-means were given the correct numbers of clusters for each dataset.

In the semi-supervised setting, we benchmarked the DPHM model against semi-supervised GMMs (Georgi et al., 2010) and semi-supervised DPGMs with diagonal covariance matrices, and constrained k-means (Wagstaff et al., 2001). Each dataset was partitioned into 10 equal sized groups, and 10-fold cross-validation was applied to calculate ARIs for each model with fixed initializations. In each fold, one group was used as the training set and the remaining nine comprised the testing set. For a given fold, we create must-link constraints to force observations, in the training set of the fold, with the same ground-truth labels to cluster together; the semi-supervised models had to cluster observations in the testing set of the fold given the must-link constraints.

Unsupervised and semi-supervised DPHM and DPGM models were run using our 'dphmix' package. Unsupervised and semi-supervised GMMs were implemented with the 'PyMix' Python package (Georgi et al., 2010). Unsupervised and constrained k-means was run using the 'COP-Kmeans' Python implementation (https://github.com/Behrouz-Babaki/COP-Kmeans) (Babaki, 2017). For the DPHM and DPGM models, prior hyperparameters were chosen to be informative so expectations of prior distributions matched the means of their corresponding features. We also set $\alpha = 1$; tuning $\alpha$ did not improve performance. Initial cluster probability vectors were randomly generated. Parameters for the GMM were initialized from randomly generated clusters. Initial centroids for k-means were randomly generated within the bounds of the features.

##### 2.2.2 Comparison to constrained k-means on biological data

We benchmarked the performance of the DPHM against constrained k-means on the Nkx2-5 dataset with differing sizes of training sets from 10 to 100 (in intervals of 10). For each training set size, we randomly sampled 5 training sets of VISTA enhancers, with replacement. For each training set, we trained a semi-supervised DPHM model and constrained k-means with a must-link constraint on the set. For both models, we refer to the cluster with the training set as the positive cluster and count the proportion of held-out VISTA enhancers (VISTA enhancers that were not in the training set) that were predicted to be in the positive cluster. We evaluate the DPHM and constrained k-means using the precision lower bound (the proportion of regions in the positive cluster, excluding the training set, that were held-out VISTA enhancers) and the sensitivity lower bound (the proportion of held-out VISTA enhancers that were in the positive cluster). These are lower bounds because regions that are not VISTA enhancers may still be active enhancers.

##### 2.2.3 Comparison to supervised classifiers on biological data

To test the DPHMs ability to predict enhancers in the same class as VISTA enhancers in a fully supervised paradigm, we compared a fully supervised DPHM to Naive Bayes (that assumes the same distributions over features as the DPHM), an elastic-net regularized SVM, random forests and AdaBoost. We labeled the 109 VISTA enhancers with 'positive' and all other 6,100 Nkx2-5 bound regions with 'negative'. Note that we are not predicting enhancer states here; we are predicting whether regions are VISTA enhancers. We split the positive set into two roughly equal sized groups and held-out one group from the training set (55 training VISTA enhancers and 54 held-out VISTA enhancers). We trained the models on the training set that contained all negatives and training VISTA enhancers. We used the models to predict labels for the training and held-out VISTA enhancers. We repeated this 10 times with 10 different random splits of the positive set to calculate sensitivities for each model on the training and held-out VISTA enhancers. In the fully supervised paradigm, the DPHM model is fit with two must-link constraints: one that forces training VISTA enhancers to cluster together and another that forces negatives to cluster together.

Naive Bayes was implemented with the 'pomegranate' Python package (Schreiber, 2017) and SVMs, random forests and AdaBoost were implemented with the 'sklearn' Python package. Parameters for the models were set to maximize sensitivities on the held-out VISTA enhancers. For Naive Bayes, both class priors were proportional to the sizes of the classes in the training set. For linear SVM, we used the 'SGDClassifier' function with loss='hinge'. Optimal values for the 'alpha' and 'l1_ratio' parameters varied across the 10 splits. We also tried non-linear kernels such as radial basis functions but the linear kernel was optimal. Random forests were implemented with the 'RandomForestClassifier' function. Optimal values for the 'n_estimators' parameter varied from 1 to 5 and optimal values for 'criterion' varied across splits. AdaBoost was implemented with the 'AdaBoostClassifier' function and the optimal value of 'n_estimators' was 2 for all splits while the optimal 'learning_rate' varied between 2 and 5.

#### 2.3 Sensitivity Analysis

We conducted sensitivity analysis to determine whether predictions, for the mouse heart dataset, are consistent across different sizes of training data. We trained a semi-supervised DPHM model to force all 109 VISTA enhancers (complete training set) to cluster together. Then, we trained semi-supervised DPHM models with must-link constraints on 10 to 100 VISTA enhancers (in intervals of 10). For each training set size, we randomly sampled 5 training sets, with replacement, from the 109 VISTA enhancers and applied a semi-supervised DPHM model for each sample. Finally, we applied an unsupervised DPHM model (without must-link constraints) on the mouse heart dataset. $\alpha$ was set to 1 for every model. For the semi-supervised models, we define predicted active enhancers (PAEs) as regions that cluster with the training set (excluding the VISTA enhancers in the training set). For the unsupervised model, we define PAEs as the regions in the cluster with the most VISTA enhancers (the VISTA enhancers are not included in the set of PAEs).

#### 2.4 GO Enrichment Analysis

To validate predicted classes, we conducted gene ontology (GO) enrichment analysis with GREATv3 (McLean et al., 2010). We defined the foreground sets as the regions of interest (all regions in a particular class) and the background set as the 6100 regions in the mouse heart dataset that did not overlap a VISTA enhancer. P-values are corrected for multiple testing using the Benjamini and Hochberg (1995) correction, and we used a false discovery rate q-value threshold of 0.05 and fold enrichment $> 2$ to define significant GO terms.

#### 2.5 Motif Enrichment Analysis

We used MotEvo (Arnold et al., 2012) to compute posterior probabilities of TFs binding to the regions in the mouse heart dataset given the position weight matrices (PWMs) for their motifs, multi-sequence alignments (MSAs) and a phylogenetic model. Nkx2-5 bound regions in mouse were resized to 700bps from their mid-points. These 700bp regions in the mouse genome were aligned to human, rhesus macaque, rat, cow and pig genomes using LiftOver (Hinrichs et al., 2006). The regions in these five species were resized to 2kb from their mid-points. Multi-sequence alignments (MSAs) were generated for the six species using MAFFTv7 (Katoh et al., 2002), an iterative refinement method. The resized aligned sequences of the mouse genome having 700bp for all the bound regions were used as the reference sequences for MotEvo [parameters: Mode=TFBS; EMprior=0; markovorderBG=0; minposterior=0.2; unknown functional elements were not used]. The phylogenetic model was determined by a nucleotide substitution matrix and an averaged phylogenetic tree for all six species. We extracted the PWMs from the JASPAR 2016 database (Mathelier et al., 2016) and found that 559 redundant PWMs had positive posterior probabilities for at least one region in the mouse heart dataset.

We applied lasso logistic regression (using the 'sklearn' Python package) to discover motifs that discriminate functional classes of enhancers from predicted inactive regions. Four regressions were trained to discriminate four classes, respectively, from a class of predicted inactive regions where posterior probabilities across the 559 motifs were used as features for each region. In each regression model, parameters were chosen to maximize the mean accuracy in 5-fold cross-validation and the coefficient of a motif quantifies its enrichment in the class of interest compared to the class of predicted inactive regions. The significance of each coefficient in each regression model was measured through a permutation test (1000 permutations). We identified enriched motifs for each class by taking the motifs with significant (permutation test $p < 0.05$) and positive coefficients for the class. For each class, we used GOrilla (Eden et al., 2009) to identify significantly ($q < 0.05$) enriched processes for TFs associated with their motifs. All TFs for the 559 motifs in the regressions were used as the background set for GOrilla.

### 3 Results

#### 3.1 Classes Found by the DPHM

Clustering the clusters allowed the DPHM model to discover five large classes of Nkx2-5 bound regions in the Nkx2-5 dataset. Here, we discuss four large classes (classes 1, 3, 4 and 5) that were not discussed in the manuscript.

The largest class (class 1) contains 2463 regions from 10 clusters, and appears to contain weak, poised, and primed enhancers (Ernst et al., 2011; Choukrallah et al., 2015) as most of the regions have high H3K4 mono-methylation (K4me1), low K27ac in embryonic mouse tissue, and are bound to P300 in adult tissue. The genes near (within 1Mb) these regions were not significantly ($q < 0.05$) enriched with any heart-specific biological processes.

Class 3 is the second largest class and contains 1176 regions across 11 clusters. This class has the highest average H3K4 tri-methylation (K4me3), H3K9 acetylation (K9ac), DNase-seq signals (in both species) and K27ac in embryonic mouse tissue; all of these features are associated with active promoters (Ernst et al., 2011). Furthermore, over 96% of regions were bound to P300 and POL2 and 794 regions overlapped with transcription start sites (TSSs) of protein-coding genes. Another 224 were within 2kb of a TSS, suggesting that they could be promoters. GREAT indicated that the promoters of their nearest genes were significantly enriched with the Yy1 binding motif ($q = 6.22 \times 10^{-11}$); Yy1 (along with other TFs) binds to active enhancers and promoters to mediate interactions between them (Weintraub et al., 2017). The nearest (or overlapping) genes were enriched with 70 biological processes, none of which were specific to heart. Instead, they were enriched with DNA conformation change ($q = 2.26 \times 10^{-10}$) and general functions such as DNA metabolic process ($q = 5.16 \times 10^{-10}$), suggesting that Nkx2-5 binds to promoters of genes that are important for chromatin remodeling and house-keeping functions.

Class 4 only has one cluster of 663 regions and appears to largely contain inactive regions (Pradeepa, 2017) with very low K4me1 and DNase-seq signals (in both species) and K27ac in embryonic mouse tissue compared to the other enhancer classes (class 1 and the DPHM-AEC). Furthermore, most regions were not bound to P300, CTCF or POL2 in adult tissue. The top two enriched biological processes were regulation of cardiac muscle cell differentiation ($q = 9.79 \times 10^{-4}$) and positive regulation of neuron differentiation ($q = 6.63 \times 10^{-3}$), suggesting that this class contains decommissioned enhancers that retained Nkx2-5 binding and played a role in heart and brain differentiation before deactivation.

Class 5 contained regions with characteristics of active enhancers along with K4me3 and K9ac, histone modifications that are typically stronger in active promoters. Ernst et al. (2011) showed that some active enhancers have these additional modifications and are usually cell-type specific. The class had 565 regions across 8 clusters and 111 of them overlapped with TSSs of protein-coding genes. An additional 251 were within 5kb of a TSS, suggesting that some may be active promoters or proximal elements. Although they were not significantly enriched with any biological processes, six genes (Hand2, Mef2c, Nkx2-5, Smarcd3, Sox4 and Tbx5) associated with cardiac ventricle formation ($q = 6.58 \times 10^{-2}$) were near regions from the class, which shows that high levels of K27ac, K4me1/2/3 and K9ac may mark heart-specific enhancers.

#### 3.2 Multinomial Lasso Logistic Regression

We repeated the motif enrichment analysis in section 2.5 with a multinomial lasso logistic regression which predicts the five classes from each other rather than predicting a functional enhancer class from predicted inactive regions. This means each class receives a vector of coefficients (one coefficient for each motif) that quantify the enrichments of the motifs in the class compared to all other classes. Similar to the results of the binomial lasso logistic regression discussed in section 3.3.2 of the manuscript, many motifs were uniquely enriched for each class (Supplementary Figure 4). In particular, 58 motifs were enriched for the DPHM-AEC and their associated TFs were significantly enriched for response to hormone ($q = 4.36 \times 10^{-2}$) and regulation of muscle system process ($4.29 \times 10^{-2}$). The other classes were not enriched for any processes.

### 4 Reproducing our Clusters

This section shows how to use the 'dphmix' package and replicate the clusters discussed in the paper.

1. First, ensure Python3 is installed and download the package with 'pip' by executing the following command in a terminal:

```
pip install dphmix
```

2. Download the Nkx2-5 dataset from https://github.com/tahmidmehdi/dphmix/tree/master/data.

3. In Python, import the package and store the data in a dataframe. The rows represent observations and columns represent features:

```
from dphmix.VariationalDPHM import *
X = pd.read_csv('nkxData.csv', index_col=0, header=0)
X.drop(['Cluster', 'Class'], axis=1, inplace=True)
```

4. Set hyperparameters for the model. Descriptions for the hyperparameters can be found on Github (https://github.com/tahmidmehdi/dphmix). We tried several different priors, including ones where the expectations of Gaussian and Bernoulli parameters were equal to the means of their corresponding features, but this prior resulted in the highest ELBO.

```
nu = [0]*100
rho = [1]*100
a = [1]*100
b = [1]*100
gamma = [1]*5
delta = [1]*5
zeta = 1/np.std(X[X.columns[105:]])
eps = zeta*np.mean(X[X.columns[105:]])
h = dict(nu=nu, rho=rho, a=a, b=b,
         gamma=gamma, delta=delta, eps=eps, zeta=zeta)
```

5. Initialize a 'VariationalDPHM' model and fit it to X. Features with 'float' data types are Gaussian. Features with 'int' data types that only have values of 0 or 1 are Bernoulli. Features with 'int' that only have non-negative values with at least one integer greater than 1 are Poisson.

```
model = VariationalDPHM(alpha=1, iterations=1000, max_clusters=100,
                        tol=10, random_state=42)
clusters = model.fit_predict(X, hyperparameters=h)
```

cluster.c will return an array of cluster indices for observations in X (the ith element is the cluster of the ith observation).

### 5 References


Arnold, P. et al. (2012). Motevo: integrated bayesian probabilistic methods for inferring regulatory sites and motifs on multiple alignments of dna sequences. Bioinformatics (Oxford, England), 28(4), 48794.

Babaki, B. (2017). Cop-kmeans version 1.5.

Benjamini, Y. and Hochberg, Y. (1995). Controlling the false discovery rate: a practical and powerful approach to multiple testing. Journal of the Royal Statistical Society. Series B (Methodological), pages 289300.

Choukrallah, M. et al. (2015). Enhancer repertoires are reshaped independently of early priming and heterochromatin dynamics during b cell differentiation. Nat Commun., 6(8324).

Eden, E. et al. (2009). Gorilla: a tool for discovery and visualization of enriched go terms in ranked gene lists. BMC Bioinformatics, 10(1), 48.

Ernst, J. et al. (2011). Mapping and analysis of chromatin state dynamics in nine human cell types. Nature, 473(7345), 4349.

Georgi, B. et al. (2010). Pymix - the python mixture package - a tool for clustering of heterogeneous biological data. BMC Bioinformatics, 11(1), 9.

Hinrichs, A. et al. (2006). The ucsc genome browser database: update 2006. Nucleic acids research, 34(suppl 1), D590D598.

Hubert, L. and Arabie, P. (1985). Comparing partitions. Journal of Classification, 2(1), 193218.

Katoh, K. et al. (2002). Mafft: a novel method for rapid multiple sequence alignment based on fast fourier transform. Nucleic Acids Research, 30(14), 30593066.

Mathelier, A. et al. (2016). Jaspar 2016: a major expansion and update of the open-access database of transcription factor binding profiles. Nucleic Acids Research, 44(D1), D110D115.

McLean, C. Y. et al. (2010). Great improves functional interpretation of cis-regulatory regions. Nature biotechnology, 28(5), 495501.

Pradeepa, M. M. (2017). Causal role of histone acetylations in enhancer function. Transcription, 8(1), 4047.

Schreiber, J. (2017). Pomegranate: fast and exible probabilistic modeling in python. Journal of Machine Learning Research, 18, 164:1-164:6.

Weintraub, A. S. et al. (2017). Yy1 is a structural regulator of enhancer-promoter loops. Cell, 171(7), 15731588.e28.

### 6 Supplementary Tables & Figures

We used the following datasets to extract features for the Nkx2-5 dataset.

| Species | Assay | Accession |
|---|---|---|
| Homo sapiens | DNase-seq–embryonic-117day | ENCFF008DNO |
| Homo sapiens | DNase-seq–embryonic-96day | ENCFF135BIN |
| Homo sapiens | DNase-seq–embryonic-116day,98day | ENCFF162WFN |
| Homo sapiens | DNase-seq–embryonic-116day,98day | ENCFF193CHS |
| Homo sapiens | DNase-seq–embryonic-103day | ENCFF281GOQ |
| Homo sapiens | DNase-seq–embryonic-96day | ENCFF326FWY |
| Homo sapiens | DNase-seq–embryonic-59day,76day | ENCFF327NVK |
| Homo sapiens | DNase-seq–embryonic-59day,76day | ENCFF351UQV |
| Homo sapiens | DNase-seq–embryonic-147day | ENCFF461COG |
| Homo sapiens | DNase-seq–embryonic-72day,76day | ENCFF498ECZ |
| Homo sapiens | DNase-seq–embryonic-110day | ENCFF509TTX |
| Homo sapiens | DNase-seq–embryonic-105day | ENCFF604EJP |
| Homo sapiens | DNase-seq–embryonic-72day,76day | ENCFF702IWR |
| Homo sapiens | DNase-seq–embryonic-80day | ENCFF751LRX |
| Homo sapiens | DNase-seq–embryonic-110day | ENCFF782WKY |
| Homo sapiens | DNase-seq–embryonic-101day | ENCFF802UAF |
| Homo sapiens | DNase-seq–embryonic-120day | ENCFF851MCI |
| Homo sapiens | DNase-seq–embryonic-91day | ENCFF854MAD |
| Homo sapiens | DNase-seq–embryonic-105day | ENCFF915OOI |
| Homo sapiens | ChIP-seq-H3K9ac-human-embryonic-105day | ENCFF064NJL |
| Homo sapiens | ChIP-seq-H3K4me1-human-embryonic-91day | ENCFF366DMT |
| Homo sapiens | ChIP-seq-H3K9me3-human-embryonic-101day | ENCFF523QJV |
| Homo sapiens | ChIP-seq-H3K4me3-human-embryonic-91day | ENCFF650XMH |
| Homo sapiens | ChIP-seq-H3K27me3-human-embryonic-101day | ENCFF665DKU |
| Homo sapiens | ChIP-seq-H3K36me3-human-embryonic-101day | ENCFF688KQB |
| Homo sapiens | ChIP-seq-H3K4me1-human-embryonic-105day | ENCFF965PJE |
| Homo sapiens | whole-genomeshotgunbisulfitesequencing–embryonic-101day | ENCFF560SMW |
| Homo sapiens | DNase-seq–embryonic-101day | ENCFF920CTO |
| Homo sapiens | DNase-seq–embryonic-136day | ENCFF185JEB |
| Homo sapiens | DNase-seq–embryonic-103day,101day | ENCFF188LTL |
| Homo sapiens | DNase-seq–embryonic-103day,101day | ENCFF791FRX |
| Homo sapiens | DNase-seq–embryonic-103day,101day | ENCFF062BSK |
| Homo sapiens | DNase-seq–embryonic-103day,101day | ENCFF581KWV |
| Mus musculus | DNase-seq–embryonic-10.50day | ENCFF071TLE |
| Mus musculus | DNase-seq–embryonic-11.50day | ENCFF503WJK |
| Mus musculus | DNase-seq–po stnatal-0day | ENCFF513QAB |
| Mus musculus | ChIP-seq-H3K4me2-mouse-embryonic-13.5day | ENCFF019ABU |
| Mus musculus | ChIP-seq-H3K27ac-mouse-embryonic-15.5day | ENCFF021RTZ |
| Mus musculus | ChIP-seq-H3K27ac-mouse-embryonic-14.5day | ENCFF034YQZ |
| Mus musculus | ChIP-seq-H3K9ac-mouse-po stnatal-0day | ENCFF042PHG |
| Mus musculus | ChIP-seq-H3K27me3-mouse-embryonic-15.5day | ENCFF044FTG |
| Mus musculus | ChIP-seq-H3K9ac-mouse-embryonic-13.5day | ENCFF045OSV |
| Mus musculus | ChIP-seq-H3K36me3-mouse-embryonic-13.5day | ENCFF046KVG |
| Mus musculus | ChIP-seq-H3K9me3-mouse-po stnatal-0day | ENCFF046SEF |
| Mus musculus | ChIP-seq-H3K27me3-mouse-embryonic-10.5day | ENCFF046VWN |
| Mus musculus | ChIP-seq-H3K4me3-mouse-embryonic-13.5day | ENCFF054NVM |
| Mus musculus | ChIP-seq-H3K4me2-mouse-embryonic-15.5day | ENCFF062SWO |
| Mus musculus | ChIP-seq-H3K4me2-mouse-po stnatal-0day | ENCFF099OBO |
| Mus musculus | ChIP-seq-H3K4me3-mouse-po stnatal-0day | ENCFF104GKL |
| Mus musculus | ChIP-seq-H3K4me3-mouse-embryonic-15.5day | ENCFF109NPC |
| Mus musculus | ChIP-seq-H3K27me3-mouse-embryonic-14.5day | ENCFF129FZH |
| Mus musculus | ChIP-seq-H3K4me1-mouse-embryonic-12.5day | ENCFF131MXX |
| Mus musculus | ChIP-seq-H3K36me3-mouse-embryonic-14.5day | ENCFF152GZF |
| Mus musculus | ChIP-seq-H3K4me2-mouse-embryonic-14.5day | ENCFF159AEV |

| | | |
|---|---|---|
| Mus musculus | ChIP-seq-H3K9me3-mouse-embryonic-10.5day | ENCFF172OCI |
| Mus musculus | ChIP-seq-H3K9me3-mouse-embryonic-11.5day | ENCFF207EPW |
| Mus musculus | ChIP-seq-H3K27ac-mouse-embryonic-10.5day | ENCFF222TEU |
| Mus musculus | ChIP-seq-H3K4me3-mouse-embryonic-14.5day | ENCFF227BHG |
| Mus musculus | ChIP-seq-H3K27me3-mouse-embryonic-16.5day | ENCFF280UTR |
| Mus musculus | ChIP-seq-H3K27ac-mouse-embryonic-11.5day | ENCFF305IHU |
| Mus musculus | ChIP-seq-H3K36me3-mouse-embryonic-15.5day | ENCFF309BMY |
| Mus musculus | ChIP-seq-H3K4me2-mouse-embryonic-11.5day | ENCFF311RNJ |
| Mus musculus | ChIP-seq-H3K4me3-mouse-embryonic-12.5day | ENCFF331OQE |
| Mus musculus | ChIP-seq-H3K27me3-mouse-embryonic-12.5day | ENCFF379MVJ |
| Mus musculus | ChIP-seq-H3K9me3-mouse-embryonic-14.5day | ENCFF409BJZ |
| Mus musculus | ChIP-seq-H3K4me1-mouse-embryonic-14.5day | ENCFF415BLI |
| Mus musculus | ChIP-seq-H3K9ac-mouse-embryonic-16.5day | ENCFF440SVM |
| Mus musculus | ChIP-seq-H3K36me3-mouse-embryonic-10.5day | ENCFF474PLN |
| Mus musculus | ChIP-seq-H3K36me3-mouse-embryonic-11.5day | ENCFF477XJO |
| Mus musculus | ChIP-seq-H3K4me1-mouse-embryonic-10.5day | ENCFF513MGM |
| Mus musculus | ChIP-seq-H3K27me3-mouse-embryonic-11.5day | ENCFF551QTH |
| Mus musculus | ChIP-seq-H3K4me3-mouse-embryonic-11.5day | ENCFF564SDZ |
| Mus musculus | ChIP-seq-H3K9ac-mouse-embryonic-12.5day | ENCFF566GYY |
| Mus musculus | ChIP-seq-H3K4me1-mouse-po stnatal-0day | ENCFF577SJR |
| Mus musculus | ChIP-seq-H3K9me3-mouse-embryonic-16.5day | ENCFF637DCW |
| Mus musculus | ChIP-seq-H3K9me3-mouse-embryonic-13.5day | ENCFF637INN |
| Mus musculus | ChIP-seq-H3K4me2-mouse-embryonic-16.5day | ENCFF652SDD |
| Mus musculus | ChIP-seq-H3K4me2-mouse-embryonic-12.5day | ENCFF671LRY |
| Mus musculus | ChIP-seq-H3K4me1-mouse-embryonic-13.5day | ENCFF673JMS |
| Mus musculus | ChIP-seq-H3K36me3-mouse-embryonic-16.5day | ENCFF686TFW |
| Mus musculus | ChIP-seq-H3K27ac-mouse-embryonic-14.5day | ENCFF689PCR |
| Mus musculus | ChIP-seq-H3K36me3-mouse-po stnatal-0day | ENCFF690EIB |
| Mus musculus | ChIP-seq-H3K9ac-mouse-embryonic-11.5day | ENCFF702AEO |
| Mus musculus | ChIP-seq-H3K4me1-mouse-embryonic-14.5day | ENCFF730VFB |
| Mus musculus | ChIP-seq-H3K4me1-mouse-embryonic-16.5day | ENCFF746ONG |
| Mus musculus | ChIP-seq-H3K9ac-mouse-embryonic-14.5day | ENCFF770FZI |
| Mus musculus | ChIP-seq-H3K4me3-mouse-embryonic-14.5day | ENCFF774GGS |
| Mus musculus | ChIP-seq-H3K9me3-mouse-embryonic-12.5day | ENCFF797OAF |
| Mus musculus | ChIP-seq-H3K27ac-mouse-embryonic-16.5day | ENCFF801FFG |
| Mus musculus | ChIP-seq-H3K9ac-mouse-embryonic-15.5day | ENCFF802SIF |
| Mus musculus | ChIP-seq-H3K27ac-mouse-po stnatal-0day | ENCFF804EYG |
| Mus musculus | ChIP-seq-H3K4me3-mouse-embryonic-16.5day | ENCFF808VAT |
| Mus musculus | ChIP-seq-H3K4me1-mouse-embryonic-15.5day | ENCFF816ESX |
| Mus musculus | ChIP-seq-H3K27me3-mouse-po stnatal-0day | ENCFF845OXU |
| Mus musculus | ChIP-seq-H3K27ac-mouse-embryonic-13.5day | ENCFF870SQY |
| Mus musculus | ChIP-seq-H3K4me3-mouse-embryonic-10.5day | ENCFF889KFS |
| Mus musculus | ChIP-seq-H3K36me3-mouse-embryonic-12.5day | ENCFF906TTD |
| Mus musculus | ChIP-seq-H3K4me1-mouse-embryonic-11.5day | ENCFF917SXQ |
| Mus musculus | ChIP-seq-H3K9me3-mouse-embryonic-15.5day | ENCFF921YZN |
| Mus musculus | ChIP-seq-H3K27ac-mouse-embryonic-12.5day | ENCFF937AXD |
| Mus musculus | ChIP-seq-H3K27me3-mouse-embryonic-13.5day | ENCFF972QDT |
| Mus musculus | ChIP-seq-CTCF-mouse-adult-8week | ENCFF447SUY |
| Mus musculus | ChIP-seq-CTCF-mouse-po stnatal-0day | ENCFF139WDG |
| Mus musculus | ChIP-seq-EP300-mouse-adult-8week | ENCFF851FMZ |
| Mus musculus | ChIP-seq-EP300-mouse-po stnatal-0day | ENCFF771SCO |
| Mus musculus | ChIP-seq-POLR2A-mouse-adult-8week | ENCFF270RGP |
| Mus musculus | whole-genomeshotgunbisulfitesequencing–embryonic-10.5day | ENCFF428AXW |
| Mus musculus | whole-genomeshotgunbisulfitesequencing–embryonic-11.5day | ENCFF107UNJ |
| Mus musculus | whole-genomeshotgunbisulfitesequencing–embryonic-12.5day | ENCFF816XTJ |
| Mus musculus | whole-genomeshotgunbisulfitesequencing–embryonic-13.5day | ENCFF114EIZ |
| Mus musculus | whole-genomeshotgunbisulfitesequencing–embryonic-14.5day | ENCFF925EPJ |
| Mus musculus | whole-genomeshotgunbisulfitesequencing–embryonic-15.5day | ENCFF951TJQ |

| | | |
|---|---|---|
| Mus musculus | whole-genomeshotgunbisulfitesequencing–embryonic-16.5day | ENCFF574RDQ |
| Mus musculus | whole-genomeshotgunbisulfitesequencing–po stnatal-0day | ENCFF467UEZ |

Table 1: ENCODE datasets used for feature extraction

| | Unsupervised | | | Semi-supervised | | |
|---|---|---|---|---|---|---|
| | 10-cluster | 25-cluster | 50-cluster | 10-cluster | 25-cluster | 50-cluster |
| DPGM | $8.11 \times 10^{-5}$ | $8.93 \times 10^{-5}$ | $9.13 \times 10^{-5}$ | $3.19 \times 10^{-5}$ | $6.43 \times 10^{-5}$ | $9.08 \times 10^{-5}$ |
| GMM | $8.11 \times 10^{-5}$ | $8.88 \times 10^{-5}$ | $8.93 \times 10^{-5}$ | $1.11 \times 10^{-3}$ | $6.43 \times 10^{-5}$ | $9.08 \times 10^{-5}$ |
| k-means | $3.83 \times 10^{-3}$ | $1.62 \times 10^{-4}$ | $1.06 \times 10^{-1}$ | 1.00 | $6.01 \times 10^{-5}$ | $6.87 \times 10^{-2}$ |

Table 2: p-values for benchmarking the DPHM against comparators (DPGM, GMM and k-means) on synthetic data. For each comparator, we conducted a Mann-Whitney U test to determine if the DPHM achieved significantly higher ARIs than the comparator on each of the 3 synthetic datasets in both unsupervised and semi-supervised settings.

| Training Set Size | p-value |
|---|---|
| 10 | $5.58 \times 10^{-3}$ |
| 20 | $5.96 \times 10^{-3}$ |
| 30 | $5.96 \times 10^{-3}$ |
| 40 | $5.58 \times 10^{-3}$ |
| 50 | $6.09 \times 10^{-3}$ |
| 60 | $6.09 \times 10^{-3}$ |
| 70 | $5.58 \times 10^{-3}$ |
| 80 | $6.09 \times 10^{-3}$ |
| 90 | $5.83 \times 10^{-3}$ |
| 100 | $7.06 \times 10^{-2}$ |

Table 3: p-values for benchmarking the DPHM against constrained k-means on the Nkx2-5 dataset. For each training set size, we randomly sampled 5 training sets and conducted a Mann-Whitney U test to determine if the DPHM achieved significantly higher PLBs than constrained k-means across training sets.

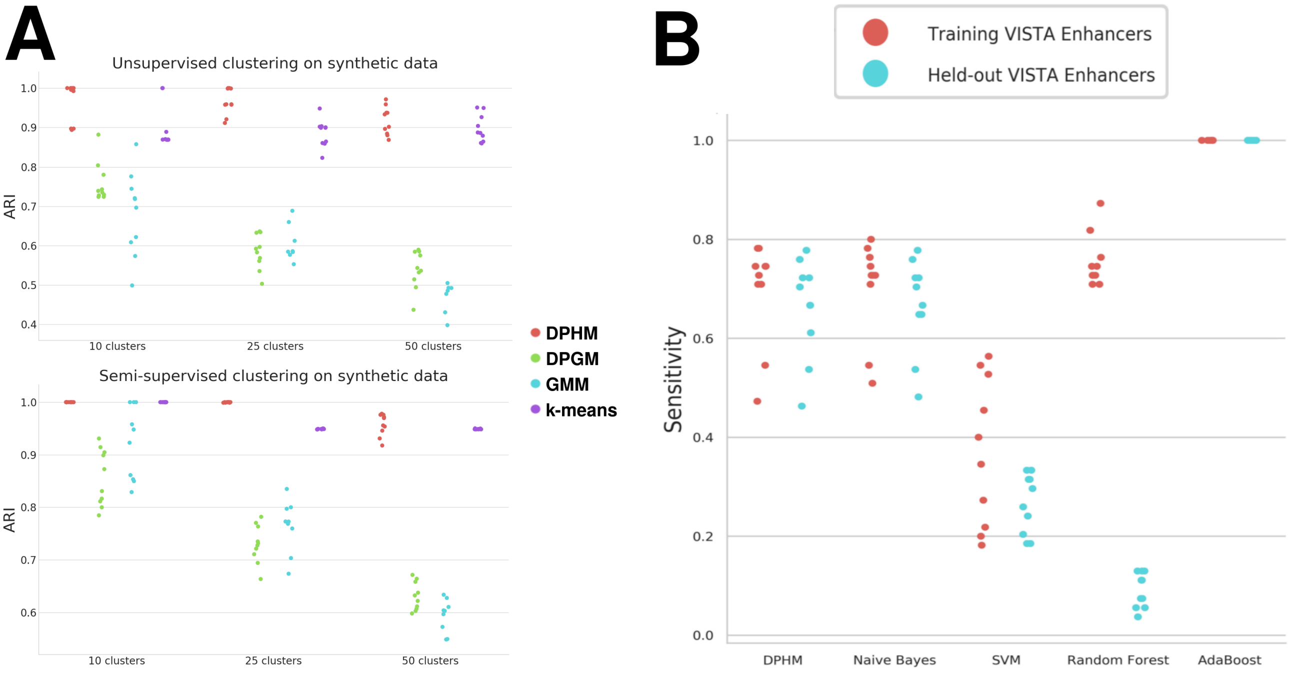


Figure 1: (A) Benchmarking results for the DPHM, DPGM, GMM and k-means on three synthetic datasets with different numbers of clusters. In the unsupervised setting, each model was applied 10 times on each dataset with different initializations and ARIs were plotted. In the semi-supervised setting, we picked the best initialization (from the unsupervised tests) and used 10-fold cross validation to compute ARIs across 10 different training sets for each model on each dataset. (B) Sensitivities of the fully supervised DPHM and supervised classifiers. Each model was applied to the Nkx2-5 dataset 10 times with 10 different random samples of the training VISTA enhancers. Sensitivities were plotted to compare their abilities to predict training and held-out VISTA enhancers.

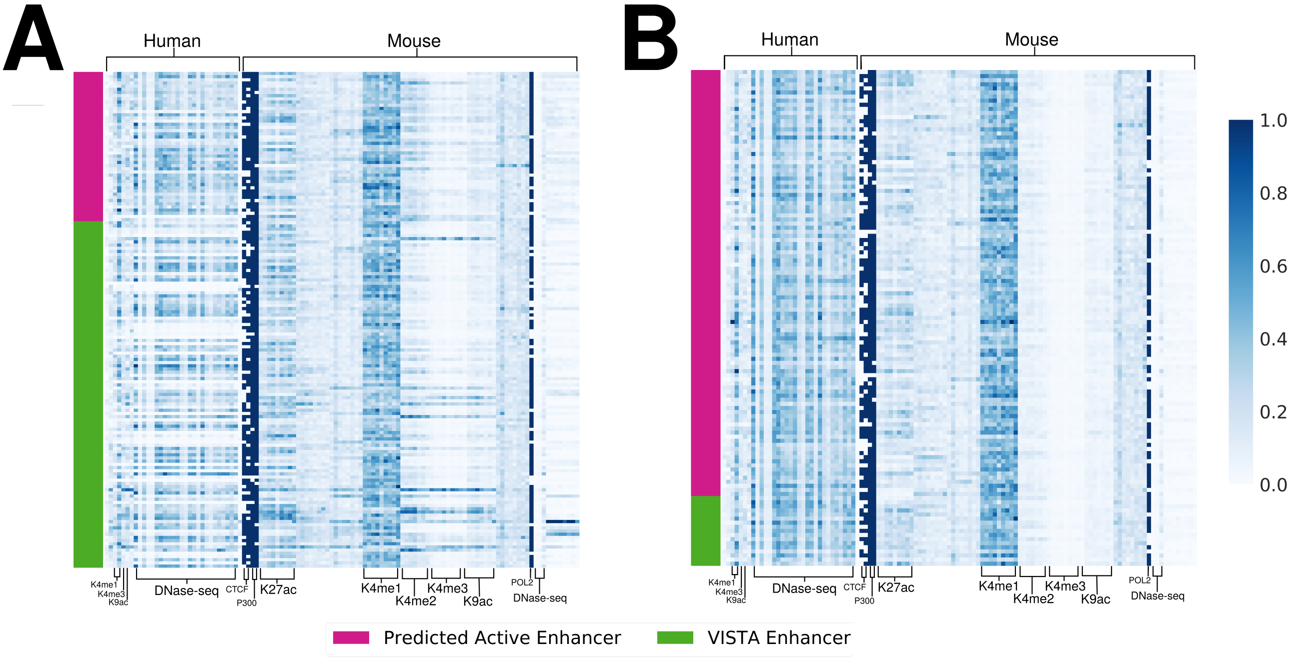


Figure 2: (A) A heatmap comparing VISTA enhancers and SDPHM-PAEs from the semi-supervised DPHM model that was trained on the complete training set. (B) A heatmap comparing VISTA enhancers and PAEs from the unsupervised DPHM model. Each column represents an ENCODE feature in heart tissues and features were minmax-scaled over the dataset to be between 0 and 1.

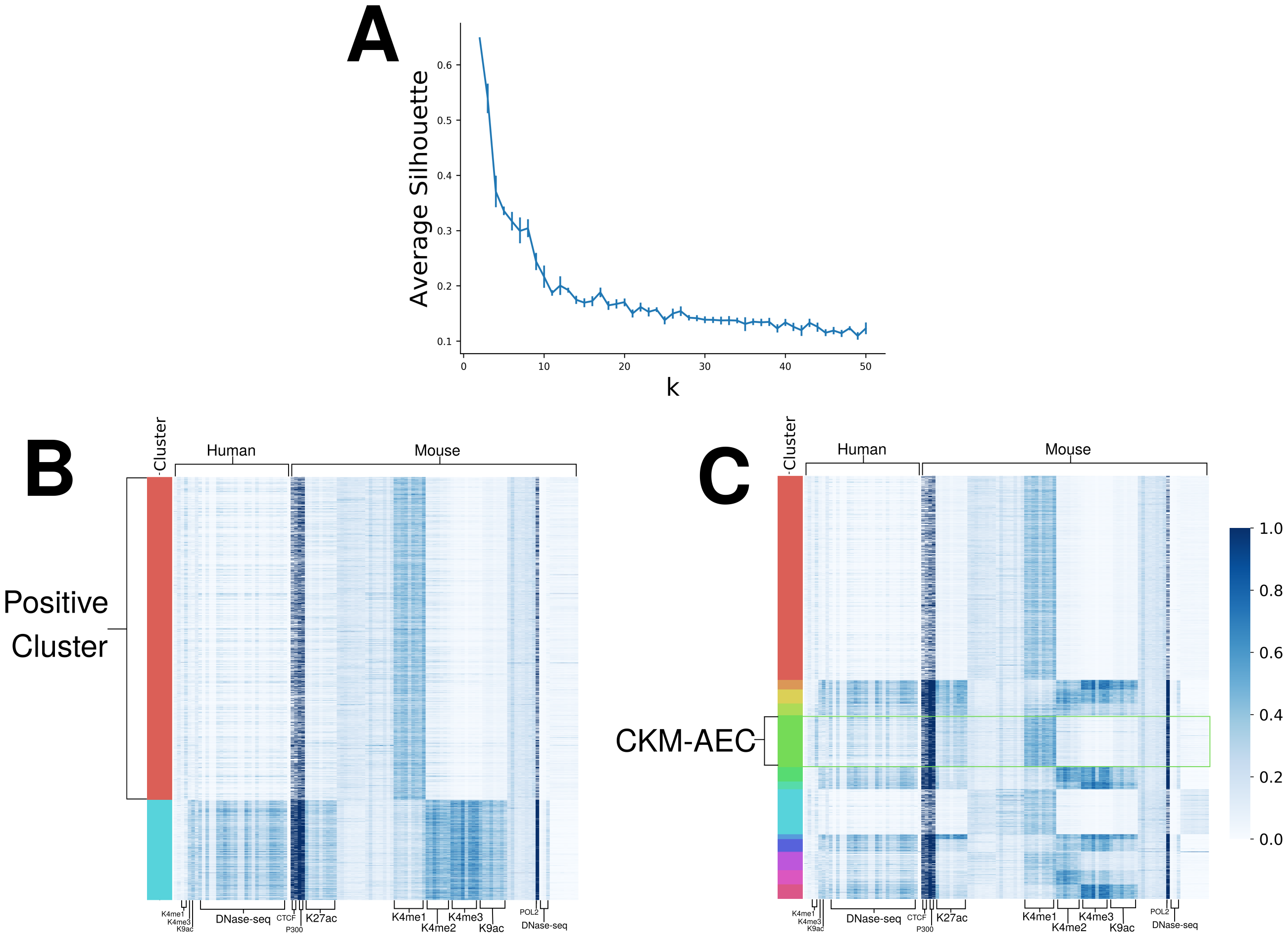


Figure 3: (A) We applied constrained k-means to the Nkx2-5 dataset with k=2 to 50 and ten different initial parameters for each k. All 109 VISTA enhancers were constrained to cluster together. The plot shows the mean average silhouette and error bars represent one standard deviation across initial parameters. (B) and (C) show heatmaps comparing clusters from constrained k-means with k=2 and k=14, respectively. Each column represents an ENCODE feature in heart tissues and features were minmax-scaled over the dataset to be between 0 and 1.

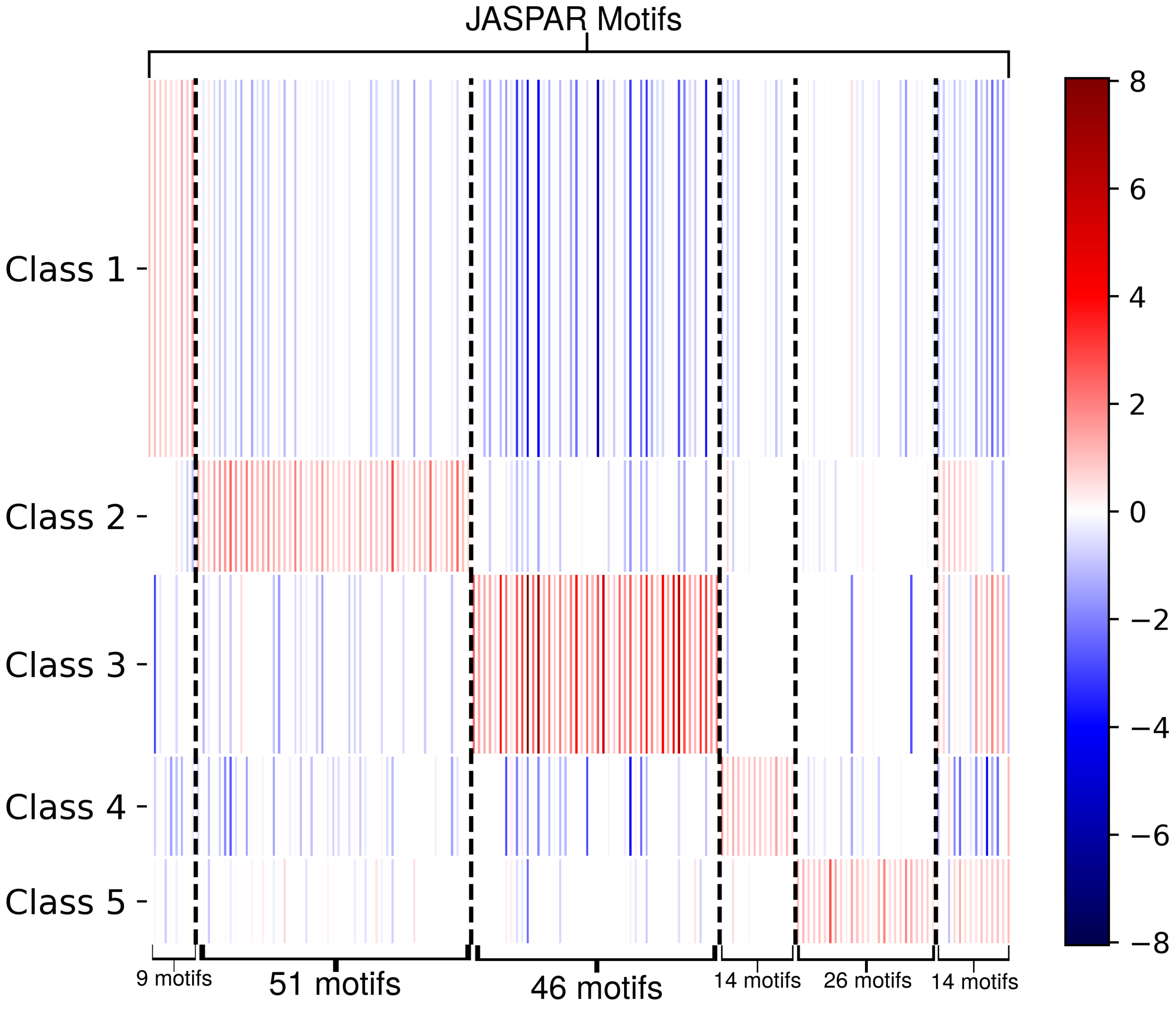


Figure 4: A heatmap showing multinomial lasso regression coefficients of JASPAR motifs in the five large classes that were found by the DPHM. The vertical sizes of each row are proportional to the number of regions in the class corresponding to the row. Only motifs that have a significant (permutation test $p < 0.05$) and positive coefficient for at least one class are included.
